# An iPSC model of fragile X syndrome reflects clinical phenotypes and reveals m^6^A- mediated epi-transcriptomic dysregulation underlying synaptic dysfunction

**DOI:** 10.1101/2024.10.14.618205

**Authors:** Lu Lu, Avijite Kumer Sarkar, Lan Dao, Yanchen Liu, Chunlong Ma, Phyo Han Thwin, Xuyao Chang, George Yoshida, Annie Li, Cenjing Wang, Crace Westerkamp, Lauren Schmitt, Maag Chelsey, Monzon Stephanie, Yu Zhao, Yaping Liu, Xiong Wang, Ling-Qiang Zhu, Dan Liu, Jason Tchieu, Makoto Miyakoshi, Haining Zhu, Christina Gross, Ernest Pedapati, Nathan Salomonis, Craig Erickson, Ziyuan Guo

**Affiliations:** Center for Stem Cell and Organoid Medicine (CuSTOM), Division of Developmental Biology, Cincinnati Children’s Hospital Medical Center, Cincinnati, OH 45229, USA; Department of Pediatrics, University of Cincinnati, College of Medicine, Cincinnati, OH 45229, USA; Division of Child and Adolescent Psychiatry, Cincinnati Children’s Hospital Medical Center, Cincinnati, OH 45229, USA; Department of Pharmacology and Toxicology, R. Ken Coit College of Pharmacy, University of Arizona, AZ 85721, USA; Division of Human Genetics, Cincinnati Children’s Hospital Medical Center, Cincinnati, OH 45229, USA; Department of Computer Science, University of Cincinnati, Cincinnati, OH 45221, USA; Department of Laboratory Medicine, Tongji Hospital, Tongji Medical College, Huazhong University of Science and Technology, Wuhan, China; Department of Pathophysiology, Key Lab of Neurological Disorder of Education Ministry, School of Basic Medicine, Tongji Medical College, Huazhong University of Science and Technology, Wuhan, Hubei, China; Division of Neurology, Cincinnati Children’s Hospital Medical Center, Cincinnati, OH 45229, USA; Division of Biomedical Informatics, Cincinnati Children’s Hospital Medical Center, Cincinnati, OH 45229, USA

## Abstract

Fragile X syndrome (FXS), the leading genetic cause of intellectual disability, arises from *FMR1* gene silencing and loss of the FMRP protein. N6-methyladenosine (m^6^A) is a prevalent mRNA modification essential for post-transcriptional regulation. FMRP is known to bind to and regulate the stability of m^6^A-containing transcripts. However, how loss of FMRP impacts on transcriptome-wide m^6^A modifications in FXS patients remains unknown. To answer this question, we generated cortical neurons differentiated from induced pluripotent stem cells (iPSC) derived from healthy subjects and FXS patients. In electrophysiology recordings, we validated that synaptic and neuronal network defects in iPSC-derived FXS neurons corresponded to the clinical EEG data of the patients from which the corresponding iPSC line was derived. In analysis of transcriptome-wide methylation, we show that FMRP deficiency led to increased translation of m^6^A writers, resulting in hypermethylation that primarily affecting synapse-associated transcripts and increased mRNA decay. Conversely, in the presence of an m^6^A writer inhibitor, synaptic defects in FXS neurons were rescued. Taken together, our findings uncover that an FMRP-dependent epi-transcriptomic mechanism contributes to FXS pathogenesis by disrupting m^6^A modifications in FXS, suggesting a promising avenue for m^6^A- targeted therapies.

## Introduction

Fragile X syndrome (FXS) is the most common inherited cause of intellectual disability, and as it often co-occurs with autism spectrum disorders (ASD), is believed to make genetic contributions to both. The syndrome manifests as a spectrum of cognitive impairments and socio-emotional deficits. FXS is primarily caused by the presence of expanded trinucleotide repeats in the fragile X messenger ribonucleoprotein 1 (*FMR1*) gene, leading to transcriptional silencing and consequent loss of Fragile X mental retardation protein (FMRP). With excessive glutamatergic signaling a pathological hallmark of FXS, previous research primarily focused on excitatory synaptic transmission, and in particular signaling downstream of the metabotropic glutamate receptor subtype 5 (mGluR5) ^1,2^. However, given the limited success of mGluR5-targeted therapies, the focus has shifted towards assessing how the balance between excitatory and inhibitory (E/I) signaling impact on network hyperexcitability ^3–6^. Despite insights gained from FMRP loss-of-function studies in animal models, clinical trials for FXS, while showing promise in preclinical studies, have yielded limited success ^7,8^. More recently, advancements have been made in modeling FXS using patient-derived induced pluripotent stem cells (iPSCs) ^9–14^, which mirror neurodevelopmental and synaptic abnormalities within the human cellular and genetic context. However, the extent to which human cultured neurons faithfully recapitulate FXS-associated disease phenotypes *in vitro* and in a way that could inform development of therapeutic strategies remains uncertain. Consequently, bridging the gap between experimental models and human disease holds paramount importance in advancing FXS research.

N6-methyladenosine (m^6^A) is a prevalent RNA modification in mammalian messenger RNAs (mRNAs). It is installed by ‘writers’ (known as methyltransferases) like the METTL3/14 complex, including METTL3, METTL14, and WTAP^15–20^, and removed by ‘erasers’ such as Fat Mass and Obesity-Associated Protein (FTO) and AlkB Homolog 5 (ALKBH5) ^20–22^. The m^6^A modification instructs regulation of RNA processes such as splicing, export, translation, and decay through the actions of m^6^A ‘readers’ that bind m^6^A-marked mRNAs^20,23–25^. In the nervous system, m^6^A modifications are particularly abundant and play established roles in brain development and neuronal functions ^26–31^. Genome-wide association studies (GWAS) have identified rare and *de novo* variants of the m^6^A reader YTHDC2 ^32–35^ that are strongly associated with ASD (SFARI Gene Score 2). However, our understanding of m^6^A regulation in human neurons, especially under pathological conditions, is still in its infancy. Consequently, transcriptome-wide m^6^A profiling in patient-derived neurons stands to significantly advance our understanding of how this modification impacts on pathological events and neuronal dysfunction.

FMRP, a multifunctional RNA-binding protein ^36^ primarily localized to translating polyribosomes, regulates local protein synthesis at synapses ^37–39^. Its deficiency results in increased synthesis of synapse-related proteins. Recently identified as a putative m^6^A ‘reader’ via unbiased proteomics studies ^40–42^, FMRP influences a range of neural processes such as neural progenitor proliferation and differentiation, synaptic transmission, plasticity, and axon growth ^43–47^. Despite a conserved consensus sequence between FMRP-binding sites and m^6^A methylation, FMRP interacts directly with m^6^A ‘readers’ in an m^6^A-modified mRNA-independent manner ^46,47^ and negatively regulates their functions ^45–47^. Specifically, FMRP antagonizes the activity of m^6^A readers, YTHDF2 and YTHDF1 (as well as the Drosophila homolog ythdf), stabilizing m^6^A- marked mRNAs and controlling their translation ^45–47^. However, the precise mechanisms by which FMRP deficiency impacts transcriptome-wide m^6^A modifications in patient-derived neurons remains elusive, and especially, to what extent this modification impacts on synaptic and neuronal dysfunction, eventually contributing to FXS pathology.

Here, we established a patient-derived iPSC neuron model of FXS and identified an imbalance in excitatory/inhibitory (E/I) signaling, along with hyperexcitability and hyperconnectivity in FXS cortical circuitry, which correlated with each individual’s clinical electroencephalogram (EEG) data. Notably, loss of FMRP leads to elevated m^6^A levels in both FXS patient-derived neurons and *Fmr1* Knockout mice brains. In mechanistic studies, we uncovered that FMRP deficiency disrupts the translational control of m^6^A writer expression, leading to accumulation of excessive m^6^A-tagged transcripts related to synapse function in FXS neurons and increased mRNA decay of such synapse-related genes. Conversely, treatment with a m^6^A writer inhibitor, capable of reducing excessive m^6^A modifications, mitigated synaptic and functional defects in FXS neurons by restoring mRNA stability. Our findings unveil a novel m^6^A-mediated epi-transcriptomic mechanism underlying FXS, by which FMRP deficiency leads to increased translation of m^6^A writers, resulting in transcriptome-wide hypermethylation and points to potential translational significance of m^6^A-modifying drugs as promising therapeutics for FXS treatment.

## Result

### Modeling FXS using patient-derived iPSCs and cortical neurons

In this study, we enrolled seven individuals with FXS and performed neuropsychological examinations on five of them, documenting significantly lower full-scale intelligence quotient (FSIQ, 45.71 ± 8, n = 5 FXS subjects), nonverbal IQ (NVIQ, 34.04 ± 8.95, n = 5 FXS subjects) and verbal IQ (VIQ, 57.38 ± 8.24, n = 5 FXS subjects), indicative of intellectual disability (Supplemental Table 1). We also validated that FMRP protein levels and FMR1 mRNA levels in blood samples from FXS patients were low or absent (Supplemental Table 1).

To model FXS, we generated induced pluripotent stem cells (iPSCs) from these seven FXS patients and age-and gender-matched healthy controls, and confirmed pluripotency, stemness, and karyotype integrity of all the iPSC lines (Supplemental Table 1). We also confirmed via PCR and Asuragen testing that CGG repeat numbers were elevated in FXS patients (over 200 CGG repeats in the 5’ UTR of the FMR1 gene compared with 30 or fewer in healthy controls; Supplemental Table 1 and Supplemental Figure 1A). The expression of forebrain cortical neural progenitor markers (Sox2, Nestin, ZO1, FOXG1, OTX2, PAX6) and mature neuronal markers (Tuj1, CaMKII), along with cortical layer markers (BRN2, CTIP2, TBR1), indicated successful induction of neuroepithelial cell fate (Figure 1A, B and Supplemental Figure 1B, C). Importantly, the FXS patient-derived cortical neurons lacked FMRP expression, unlike their control counterparts (Figure 1C, D), replicating a hallmark of FXS pathology.

**Figure 1.**
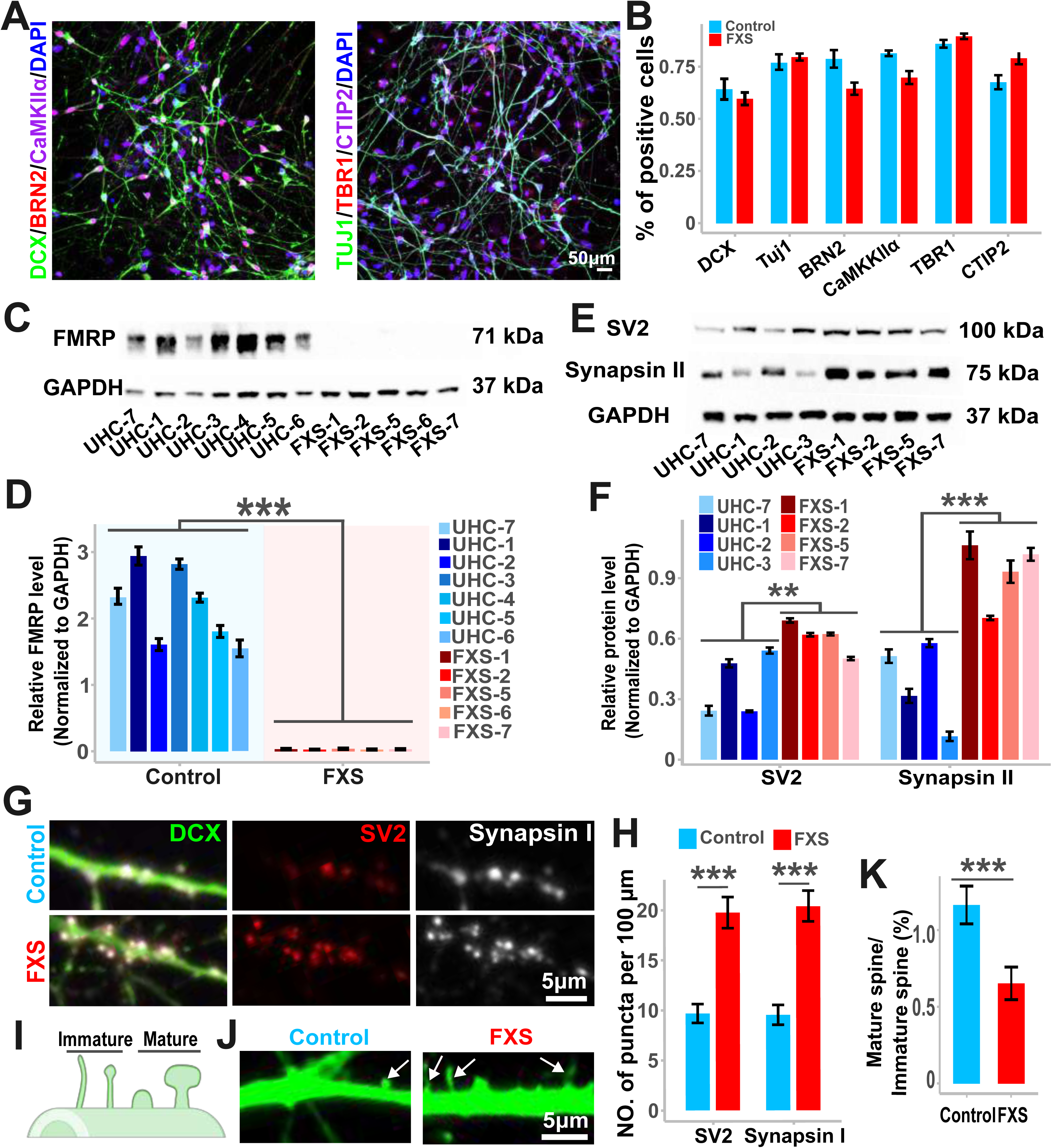
Synaptic malformations in FXS iPSC-derived cortical neurons. **A**. Confocal images of iPSC-derived neurons expressing neuronal markers DCX and TUJ1, cortical layer markers BRN2, CTIP2, and TBR1, and the glutamatergic neuron marker CaMKII. The scale bar is 50 μm. **B**. Quantification of neuronal, cortical layer, and glutamatergic marker expression. Values represent mean ± s.e.m (n = 3 biological repeats per condition). **C-D**. Western blot analysis and quantification of FMRP expression in iPSC-derived cortical neurons from unaffected healthy controls (UHC) and FXS subjects. Values represent mean ± s.e.m. (n = 3 to 5 repeats per line; ***p < 0.001, two-tailed t-test). **E-F**. Western blot analysis and quantification of synaptic proteins SV2 and Synapsin II expression in six-week-old UHC and FXS neurons. Values represent mean ± s.e.m. (n = 3 to 5 repeats per line; **p < 0.01, ***p < 0.001, two-tailed t-test). **G-H**. Representative images and quantification of synaptic puncta in six-week-old UHC and FXS neurons stained for the synaptic markers SV2 and Synapsin II. The scale bar is 5 μm. Values represent mean ± s.e.m. (n = 13 to 20 neurons per condition; ***p < 0.001, two-tailed t-test). **I-K**. Illustration of mature and immature spines, confocal images, and quantification showing an increase in the ratio of immature to mature spines in FXS iPSC-derived neurons compared to controls. Arrows point to spines. The scale bar is 5 μm. Values represent mean ± s.e.m. (n = 40 to 45 neurons per condition; ***p < 0.001, two-tailed t-test).

In western blot and immunostaining analysis, we identified elevated levels of synaptic proteins Synapsin II and SV2 (Figure 1E, F) and an increase in numbers of synaptic puncta (Figure 1G, H) in the FXS cortical neurons. We also detected distinct alterations in spine morphology, including reduced mushroom-like spines and increased numbers of filopodia-like spines compared to control neurons (Figure 1I-K). However, we did not detect any significant differences in neurites including number of sections, total neurite length, terminal points, branch points, maximum section length, minimum section length, and mean section length between FXS neurons and controls (Supplemental Figure 1D-L). Taken together, these observations align with previous research linking the absence of FMRP to changes in synaptic protein expression and synaptic structure in FXS neurons ^10,11,48–51^.

### Dysregulated synaptic transmission, excitatory/inhibitory imbalance, and hyperexcitability in cortical neurons derived from FXS iPSCs

To delve into the functional aspects of our iPSC model system, we conducted comprehensive electrophysiological analyses, including whole-cell patch-clamp and micro-electrode arrays (MEA). In whole-cell patch-clamp recordings (Figure 2A), we noted a consistent and significant increase in excitatory synaptic transmission in FXS cortical neurons relative to controls (Figure 2B). Specifically, the inter-event interval was decreased in excitatory postsynaptic currents (EPSC) whereas the frequency was elevated but the amplitude reduced (Figure 2C, D). We also observed an overall decrease in inhibitory synaptic transmission in FXS cortical neurons relative to controls (Figure 2E), manifested as an increase in inter-event interval and lower frequency and amplitudes of inhibitory postsynaptic currents (IPSC) (Figure 2F, G).

**Figure 2.**
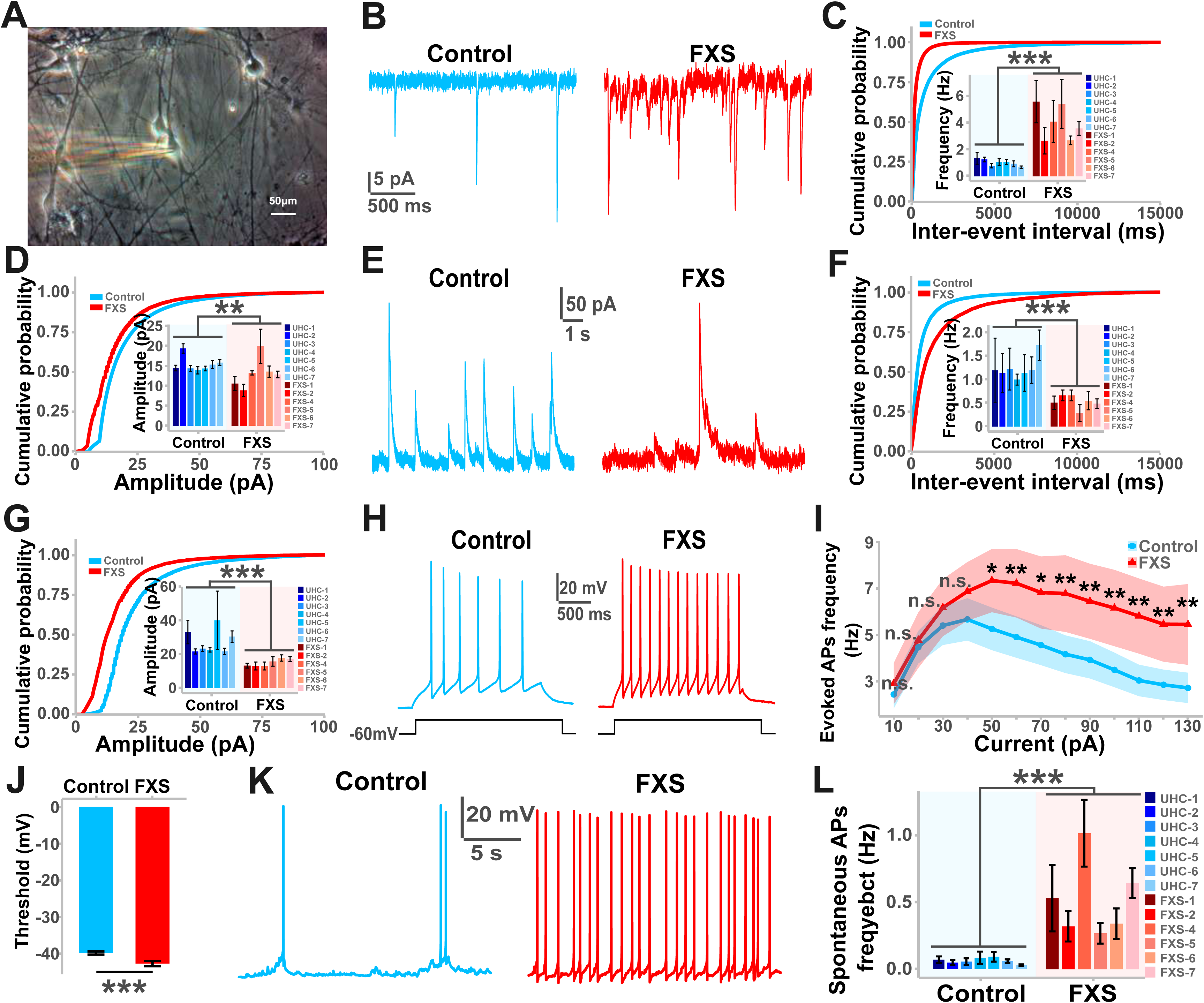
Aberrant synaptic transmission, E/I imbalance, and hyperexcitability in FXS iPSC-derived cortical neurons. **A**. Phase image showing a six-week-old FXS iPSC-derived neuron recorded via patch clamp. The scale bar is 50 μm. **B-D**. Defects in excitatory synaptic transmission in FXS iPSC-derived cortical neurons compared to control neurons. Shown are sample whole-cell voltage-clamp recording traces of excitatory postsynaptic currents (EPSCs) from six-week-old neurons (**B**) and cumulative distribution plots of spontaneous EPSC intervals (**C**, Kolmogorov–Smirnov test, the p-value is approaching zero.) and amplitudes (**D**, Kolmogorov–Smirnov test, p < 2.2e-16). Insets display quantifications of mean frequencies and amplitudes of spontaneous EPSCs. Values represent mean ± s.e.m. (n = 48 to 125 neurons per condition; **p < 0.01, ***p < 0.001, two-tailed t-test). **E-G**. Defects in inhibitory synaptic transmission in FXS iPSC-derived cortical neurons compared to control neurons. Shown are sample whole-cell voltage-clamp traces of spontaneous inhibitory postsynaptic currents (IPSCs) from six-week-old neurons (**E**, V_m_ = 20 mV), with cumulative distribution plots of spontaneous IPSC intervals (**F**, Kolmogorov–Smirnov test, the p-value is approaching zero.) and amplitudes (**G**, Kolmogorov–Smirnov test, the p-value is approaching zero.). Quantifications of mean frequencies and amplitudes of spontaneous IPSCs are also shown. Values represent mean ± s.e.m. (n = 40 to 58 neurons per condition; ***p < 0.001, two-tailed t-test). **H-L**. Increased excitability in FXS iPSC-derived cortical neurons compared to control neurons. Sample whole-cell current-clamp traces of evoked action potentials (APs) after a 60-pA injection in six-week-old neurons (**H**), with quantifications of evoked AP frequency against injected current (**I**), and threshold (**J**). Sample traces of spontaneous APs (**K**) and quantification of spontaneous AP frequency (**L**) are also provided. Values represent mean ± s.e.m. (n = 45 to 49 neurons per condition; *p < 0.05, **p < 0.01, ***p < 0.001, n.s. denotes non-significant; two-tailed t-test for threshold and spontaneous AP frequency, two-way ANOVA with Tukey’s multiple comparisons for evoked AP frequency against injected current).

Beyond the synaptic transmission defects and E/I imbalance, FXS cortical neurons were hyperexcitable, reflected as an increased frequency of evoked action potentials (APs) and decreased thresholds to firing APs (Figure 2I, J). Other parameters, including evoked AP amplitude and Rheobase, remained unchanged (Supplemental Figure 2A, B). Additionally, the frequency of spontaneous AP was higher in FXS cortical neurons relative to control cortical neurons (Figure 2K, L). However, we did not observe any significant disparities in sodium and potassium channel activities between FXS and control neurons (Supplemental Figure 2C-E). Taken together, our data thus far suggested a causal relationship between E/I imbalance and hyperexcitability in FXS iPSC-derived cortical neurons.

### Hyperactivity and hyperconnectivity in cortical circuitry derived from FXS iPSCs

To monitor cortical circuitry activity, we cultured 0.5 million neurons on a MEA chip with 64 electrodes and recorded field potentials for 5 minutes at 4, 5, 6, 7, and 8 weeks (Figure 3A and Supplemental Figure 3A). Data were sampled at 20 kHz with a 1-1000 Hz frequency bandwidth (see Methods). We identified spikes based on a threshold (5.5 times the standard deviation of peak-to-peak noise), and bursts as sequences of at least 3 spikes with inter-spike intervals within 300 milliseconds and a duration exceeding 200 milliseconds. In FXS iPSC-derived cortical circuitry, we found heightened spike and burst activity in sample traces (Figure 3B), characterized by a decreased inter-spike interval, increased burst duration, and more spikes per burst, indicative of hyperactivity (Figure 3C-E).

**Figure 3.**
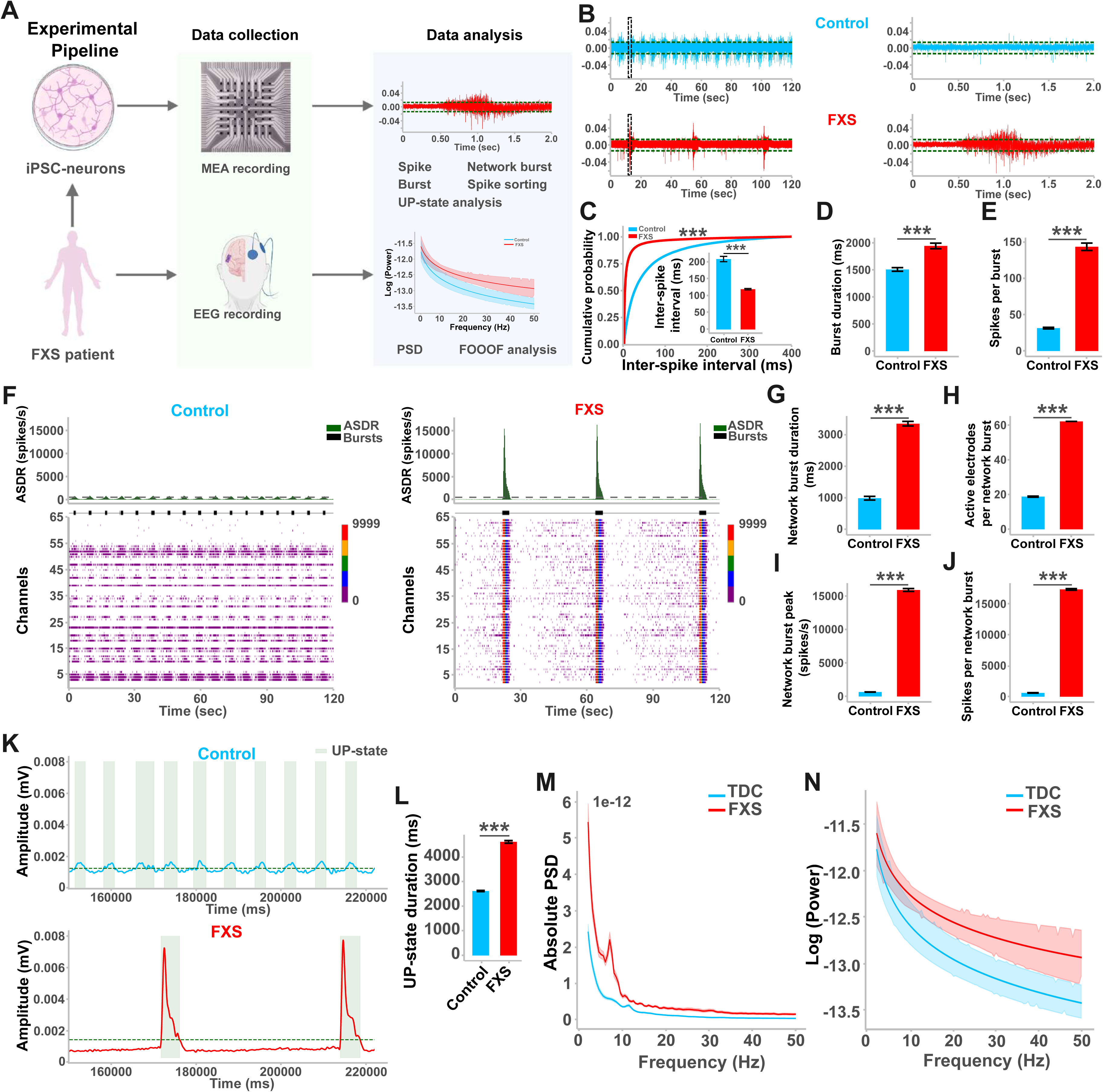
Hyperexcitability and hyperconnectivity conserved between FXS iPSC model and the same FXS subject. **A**. Schematic illustrating electrophysiological workflows for MEA and EEG recordings in both FXS subjects and their derived neurons. **B-E**. Increased excitability in FXS iPSC-derived cortical neurons compared to control neurons. Shown are sample MEA recording traces of spontaneous spikes from six-week-old neurons (**B**; right panel is an enlarged view of dashed lines in the left panel) and cumulative distribution plots of spontaneous spike intervals (**C**, Kolmogorov–Smirnov test, the p-value is approaching zero.). Quantifications of mean inter-spike interval (Inset), burst duration (**D**), and spikes per burst (**E**) are provided. Values represent mean ± s.e.m. (n = 3 cultures; ***p < 0.001, two-tailed t-test). **F-J**. Enhanced network synchrony and hyperconnectivity in FXS iPSC-derived cortical circuitry compared to controls. **F**. Sample traces and raster plots show array-wide spike detection rate (ASDR) across 64 electrodes over 120 seconds. Grey lines indicate network burst detection thresholds, black boxes indicate burst duration, and color codes represent spike frequency. Quantifications of network burst duration (**G**), active electrodes per network burst (**H**), network burst peak (**I**), and spikes per network burst (**J**) are also shown. Values represent mean ± s.e.m. (n = 3 cultures; ***p < 0.001, two-tailed t-test). **K-L**. Prolonged depolarization in FXS iPSC-derived cortical circuitry compared to controls. Shown are sample traces of MEA recording sum amplitude over 64 electrodes across 70 seconds in control and FXS cortical circuitry (**K**). Dashed lines indicate thresholds for UP-state detection, with boxes representing UP-state duration. Quantification of UP-state duration is also shown (**L**). Values represent mean ± s.e.m. (n = 3 cultures; ***p < 0.001, two-tailed t-test). **M-N**. Cortical hyperactivity and hyperconnectivity in FXS patients. Quantification of absolute power spectral density (PSD) across different frequency bands from EEG data of four typically developing control (TDC) and FXS subjects (**M**). FOOOF (Fitting Oscillations and One-Over-F) plot shows a steeper slope in FXS patients compared to TDC subjects (**N**).

Additionally, analysis of array-wide spike detection rate (ASDR) revealed enhanced neuronal network synchrony and hyperconnectivity in FXS iPSC-derived cortical microcircuitry (Figure 3F), reflected as increased burst peak and duration, more active electrodes per network burst, and elevated spike numbers in network bursts (Figure 3G-J). When sorting spikes for each electrode based on waveforms, we found significantly more waveforms in FXS iPSC-derived cortical microcircuitry (Supplemental Figure 3B, C), indicating more synchronized neurons in FXS cortical circuitry. Lastly, UP-state analysis, associated with synchronized activity across a network of neurons, particularly in the cortex, exhibited longer durations in FXS iPSC-derived cortical circuitry (Figure 3K, L). Taken together, these electrophysiological findings identify both cortical network hyperactivity and hyperconnectivity in the FXS cortical microcircuitry.

To bridge the gap between *in vitro* models and patients, we examined electroencephalogram (EEG) recordings from the same individuals and correlated these with clinical neuropsychological symptoms. We had already noted deviations in FSIQ, NVIQ, and VIQ in FXS subjects as indications of intellectual disabilities (Supplemental Table 1).

Additionally, we assessed the absolute neural power spectral densities (PSDs) of typically developing control (TDC) and FXS EEG data (Supplemental Figure 3D). We found high gamma power in FXS subjects (Figure 3M), which is associated with hyperactivity and inversely correlated with NVIQ in FXS subjects ^52–55^. Using FOOOF (Fitting Oscillations and One-Over-F) analysis to parameterize the PSDs ^56,57^, we found that the FOOOF slope was higher in FXS subjects (Figure 3N), an additional indication of cortical hyperactivity and hyperconnectivity.

Taken together, these findings strongly suggest that key pathophysiological signatures in FXS patients such as cortical hyperactivity and hyperconnectivity can be modeled in patient-derived neurons, underscoring the translational potential and clinical relevance of our human model system in uncovering disease-related mechanisms.

### FMRP-bound mRNAs encoding m^6^A writers and FMRP deficiency increasing writer expression

Like FMRP, the mRNA modification m^6^A is highly enriched in the brain, where it plays a crucial role in neurodevelopment and in mature neurons ^26–31^. To quantify total m^6^A levels, we purified mRNAs and conducted dot plot analysis using an anti-m^6^A antibody to detect m^6^A tag-tagged mRNAs. Relative to control, m^6^A levels were notably increased in FXS iPSC-derived cortical neurons (Figure 4A). Additionally, we found increased protein, but not mRNA, expression of m^6^A writers, including METTL3, METTL14, and WTAP, the key METTL3/14 complex components (Figure 4B-D). We documented similar findings in *Fmr1* knockout mice (Supplemental Figure 4). These findings suggest that FMRP deficiency leads to elevated m^6^A levels through upregulation of m^6^A writer protein expression.

**Figure 4.**
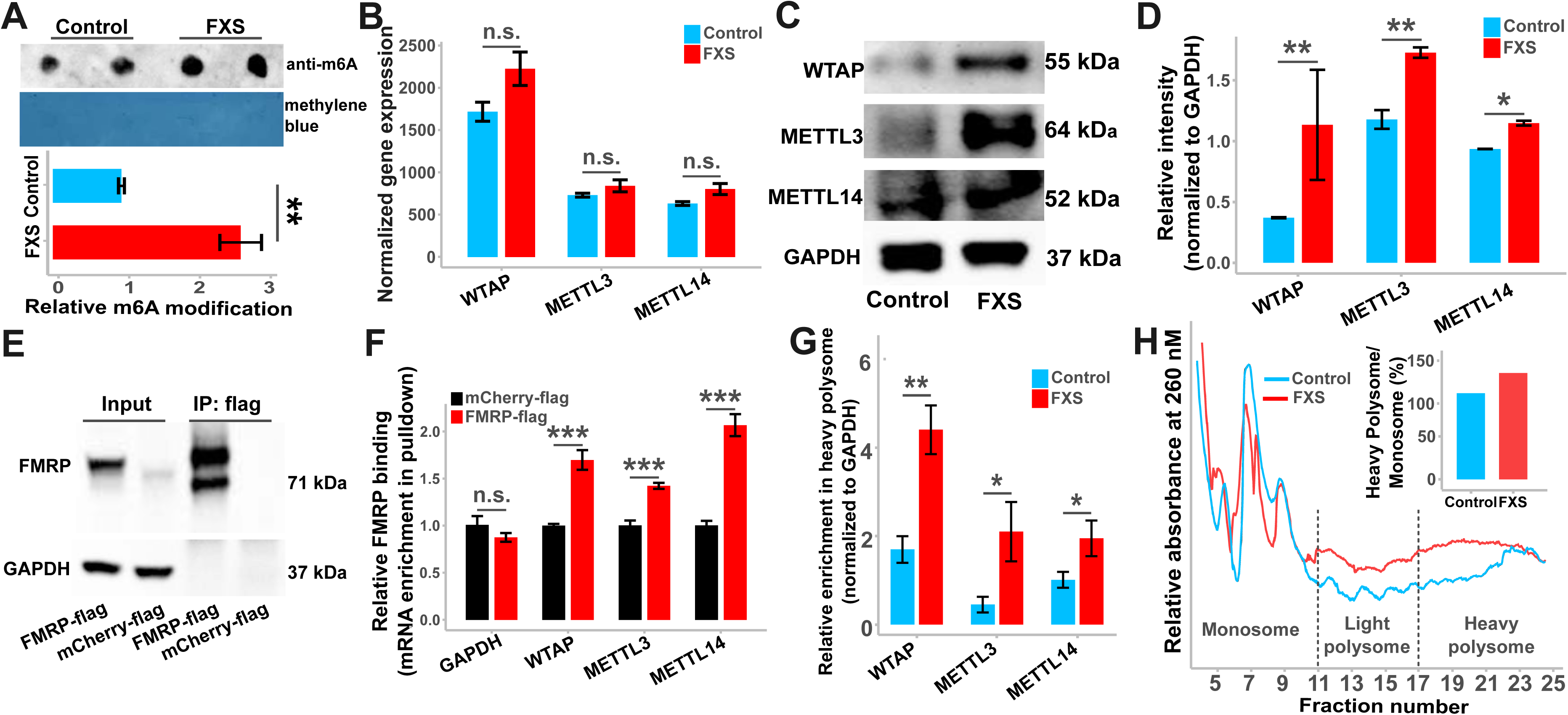
FMRP binding to mRNAs encoding m^6^A writers and the impact of FMRP deficiency on writer expression. **A**. Elevated m^6^A levels in FXS iPSC-derived cortical neurons compared to controls. Shown are dot blot assay results for polyA^+^ RNA m^6^A levels, with quantification of m^6^A intensity in FXS iPSC-derived neurons relative to controls, normalized to methylene blue. Values represent mean ± s.e.m. (n = 4 cultures; **p < 0.01; two-tailed t-test). **B**. Transcriptional expression analysis revealing no significant change of METTL3, METTL14, and WTAP transcripts between control and FXS iPSCs-derived neurons. Values represent mean ± s.e.m. (n = 7 control and 6 FXS samples; n.s. denotes non-significant; two-tailed t-test). **C-D**. Western blot analysis and quantification of METTL3, METTL14, and WTAP protein levels. Values represent mean ± s.e.m. (n = 4 cultures; *p < 0.05, **p < 0.01; two-tailed t-test). **E-F**. FMRP binds to mRNAs encoding m^6^A writers in HEK293T cells. Western blot of flag immunoprecipitation with flag-tagged mCherry–FMRP and flag-tagged mCherry, along with their input samples (**E**). Detection was performed with FMRP and GAPDH antibodies. RT-PCR quantification of GAPDH, METTL3, METTL14, and WTAP mRNA enrichment in pulldown samples (**F**) shows FMRP binding to METTL3, METTL14, and WTAP but not GAPDH mRNAs. Values represent mean ± s.e.m. (n = three biological replicates; ***p < 0.001, n.s. denotes non-significant; two-tailed t-test). **G**. RT-PCR quantification of METTL3, METTL14, and WTAP mRNAs in translationally active heavy polysome fractions from FXS iPSC-derived neurons versus controls. Values represent mean ± s.e.m. (n = 3 biological replicates; *p < 0.05, **p < 0.01; two-tailed test). **H**. Polysome profiles demonstrated a profound increase of global translation, as indicated with a 23.5% increase in the ratio of heavy polysomes to monosomes fractions (insert).

To determine whether FMRP binds mRNAs encoding m^6^A writers, we transfected HEK293T cells with flag-mCherry-FMRP and flag-mCherry plasmids, and conducted pulldown using anti-flag M2 affinity gel (Figure 4E) ^58^. In the pulldowns with flag-tagged mCherry-FMRP, but not with flag-tagged mCherry, we observed significant enrichment of METTL3, METTL14, and WTAP mRNAs (Figure 4F). There was no specific enrichment for GAPDH mRNAs, indicating that FMRP binds METTL3, METTL14, and WTAP mRNAs. To investigate whether FMRP regulates translation of the m^6^A writer translation, we analyzed the translationally active heavy polysome fractions (see Methods)^59^. Indeed, in FXS patient-derived neurons, METTL3, METTL14, and WTAP were all significantly enriched in the heavy polysome fraction (Figure 4G). In addition, the overall protein translation efficiency increased by 23.5% in FXS neurons (Figure 4H). These findings suggest that basal translation of m^6^A writer mRNAs is enhanced in the absence of FMRP.

### Transcriptome-wide m^6^A dysregulation in cortical neurons derived from FXS iPSCs

As a comprehensive profiling of transcriptome-wide m^6^A modifications in the context of FXS remains outstanding, we next performed methylated RNA immunoprecipitation sequencing (MeRIP-Seq) of neurons derived from six FXS patients and seven control samples (see Methods). When comparing datasets, we identified 3,313 m^6^A-enriched genes in FXS patient-derived neurons and 2,815 enriched genes in controls, with a widespread distribution across all chromosomes (Figure 5A). This data set indicates a transcriptome-wide neuronal dysregulation of m^6^A modifications in patients with FXS. Motif analysis and the distribution of m^6^A peaks confirmed the robustness of MeRIP-Seq and m^6^A analysis, with a predominant GGAC motif near the stop codon (Figure 5B, C). For benchmarking, we conducted MeRIP-Seq on postmortem cortex tissues derived from FXS subjects and healthy controls (Supplemental Table 2), and validated motif analysis and m^6^A peak distributions, (Supplemental Figure 5A-C).

**Figure 5.**
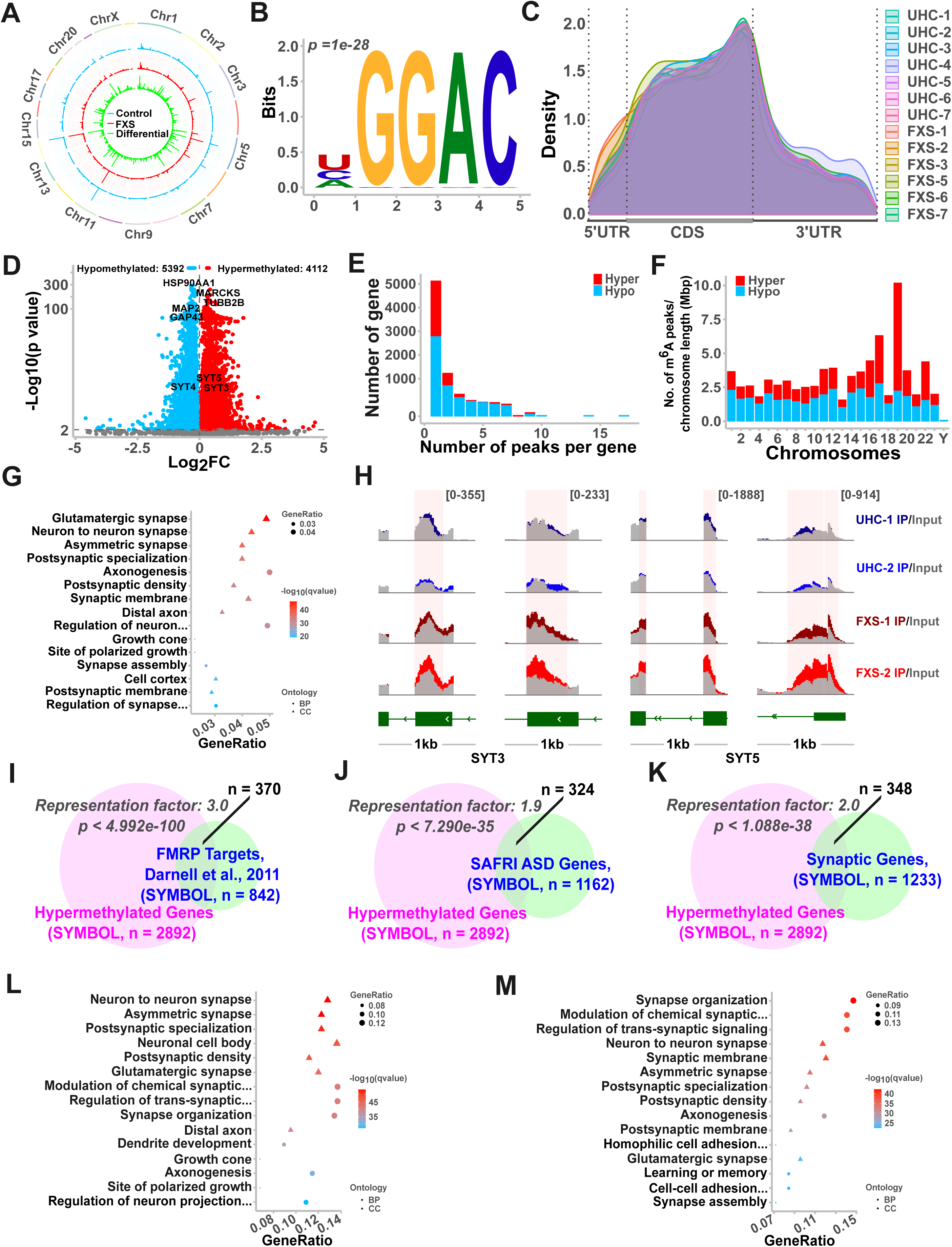
**Transcriptome-wide m^6^A dysregulation in FXS iPSC-derived cortical neurons. A-C**. Transcriptome-wide profiling of m^6^A in 6-week-old neurons from healthy controls and FXS patients. **A**. Metagene plot showing the distribution of m^6^A peaks across human chromosomes. **B**. Sequence logo illustrating the top enriched motif in m^6^A peaks (p-value = 10⁻²⁸). **C**. Distribution of m^6^A methylation along transcripts, divided into 5′ UTR, CDS, and 3′ UTR. **D**. Volcano plot depicting MeRIP-seq data for m^6^A peaks differentially methylated between FXS and control groups (n = 7 healthy controls, 6 FXS samples; adjusted p-value < 0.05; blue and red dots represent significant hypo-and hyper-methylated sites). **E-F**. Histogram showing the distribution of hyper-and hypo-methylated sites per gene (**E**) and per chromosome length across human chromosomes (**F**; blue and red boxes indicate hypo-and hyper-methylated sites). **G**. Gene Ontology (GO) analysis of differentially methylated genes (DMGs) between FXS iPSC-derived neurons and controls, revealing the top 15 enriched cellular component (CC) and biological process (BP) terms. The color represents the adjusted p-value, and the size represents the gene ratio (percentage of DMGs in each GO term). **H**. Integrative Genomics Viewer (IGV) tracks displaying MeRIP-seq read distribution for SYT3 and SYT5 mRNAs. **I-K**. Venn diagrams showing overlap between hypermethylated transcripts and FMRP target transcripts (**I**), SFARI ASD genes (**J**), and reported synapse-related genes (**K**), along with representation factors and p-values. **L-M**. GO term analysis of genes overlapping between hypermethylated transcripts and FMRP targets (**L**), as well as genes overlapping with SAFRI ASD genes (**M**).

Notably, we identified over 9,000 dys-methylated sites between controls and FXS neurons (Figure 5D) and over 4,000 dys-methylated sites between control and FXS postmortem cortex tissues (Supplemental Figure 5D), indicating transcriptome-wide m^6^A dysregulation in cultured FXS neurons and in brain tissues. Interestingly, the majority of genes exhibited one hyper-or hypo-methylation site across all chromosomes (Figure 5 E, F and Supplemental Figure 5E, F).

When we conducted gene ontology (GO) analysis, we identified significant enrichment of genes with elevated m^6^A modifications in categories related to synaptic assembly, structure, and function (Figure 5G and Supplemental Figure 5G). Examples include hypermethylation of genes encoding synaptic proteins, such as SYT3 and SYT5, in exons near the stop codon and 3’ UTR (Figure 5H). Notably, genes with elevated m^6^A modifications overlapped significantly with previously identified FMRP target genes ^38^, autism-associated genes from the SFARI database, and synapse-related genes from the SynGO database (Figure 5I-K and Supplemental Figure 5H-J). These data underscore the critical role of excessive m^6^A tagging in FXS pathologies, particularly synaptic defects, and its broader implications for autism-spectrum disorders.

Furthermore, the prominent overlap between hypermethylated genes and FMRP targets, as well as autism-associated genes, was strongly linked to synapse assembly and function (Figure 5L, M and Supplemental Figure 5K, L), reinforcing that dysregulated m^6^A modifications and synaptic dysfunction are central to FXS and autism-spectrum disorders.

### Dysregulated synapse-related gene expression

We next performed RNA sequencing (RNA-seq) on iPSC-derived neurons from seven healthy controls and six FXS patients (see Methods) and identified over 2,000 significantly differentially expressed genes (DEGs) (Figure 6A). Among these, FMR1 was prominently silenced in FXS iPSC-derived neurons (Figure 6B), consistent with full mutation status (Supplemental Table 1) and loss of FMRP expression (Figure 1C, D). Through functional enrichment analysis, we identified that nearly all top DEGs categories were linked to synaptic structure and function (Figure 6C). Moreover, DEGs significantly overlapped with FMRP target genes, particularly those associated with synapse-related categories and previously reported synapse-related genes (Supplemental Figure 6A-C). These findings highlight gene expression alterations driving synaptic defects in our iPSC-derived cortical neuron model. Similarly, RNA-seq analysis of cortical postmortem tissues from three FXS patients and healthy controls identified nearly 1,500 significantly altered genes (Supplemental Figure 6D). These genes overlapped with FMRP target and autism-associated genes from the SFARI database (Supplemental Figure 6E-I), highlighting that dysregulated synapse-related gene expression serves as a molecular signature of FXS.

**Figure 6.**
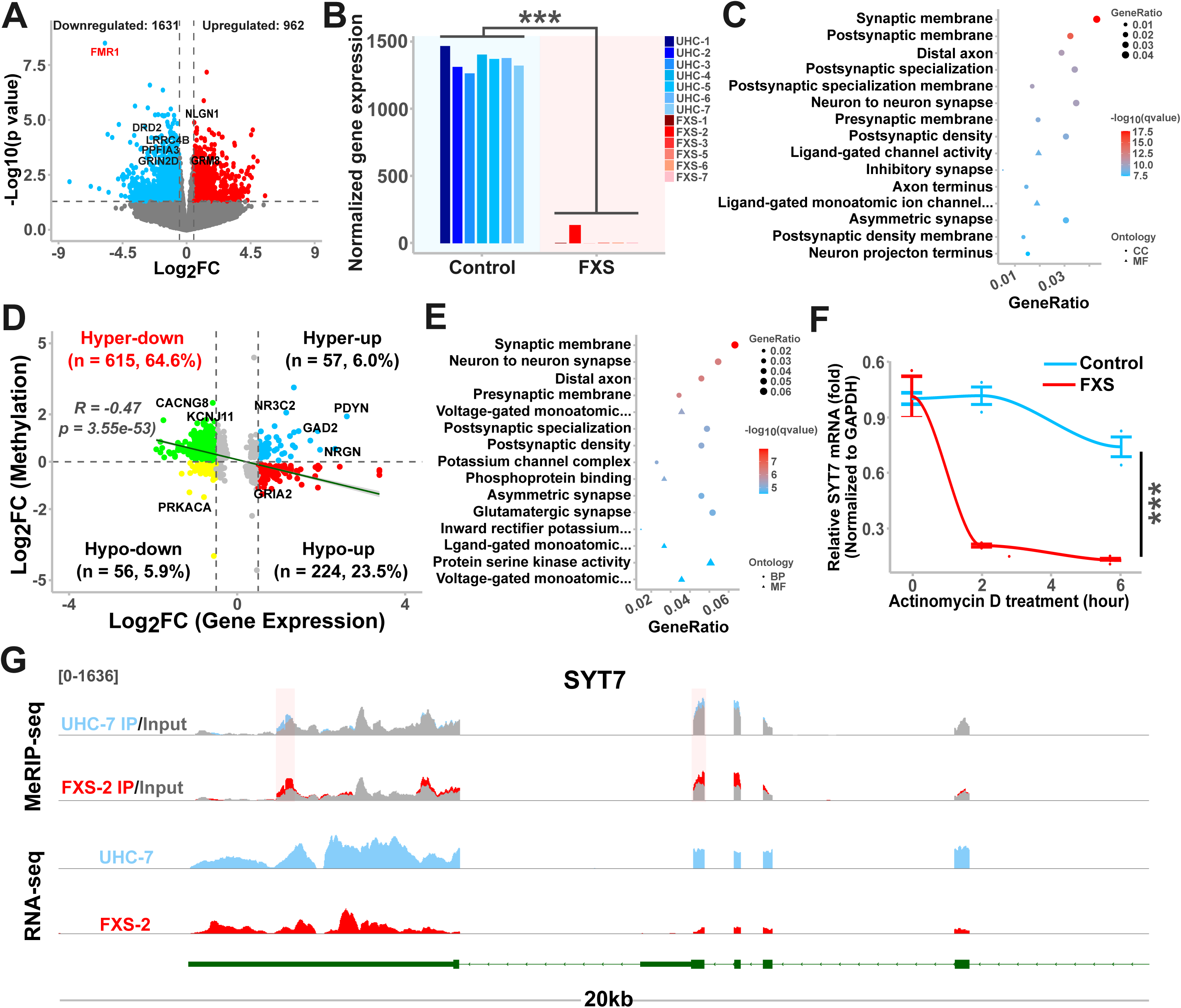
Correlation of hypermethylation with gene downregulation and increased mRNA decay in synapse-related genes. **A**. Volcano plot displaying differential gene expression between FXS iPSC-derived neurons and controls (n = 7 healthy controls, 6 FXS samples; fold change > 1.4, p-value < 0.05; blue and red dots represent significantly down-and up-regulated genes, respectively). **B**. Transcriptional expression analysis revealing significant abundant FMR1 transcripts in control but not FXS iPSCs-derived neurons. Values are shown as mean ± s.e.m. (n = 7 control, 6 FXS samples; *** p < 0.001; two-tailed t-test). **C**. Gene Ontology (GO) term analysis of differentially expressed genes (DEGs) between FXS iPSC-derived neurons and controls, showing the top 15 enrichment terms for cellular component (CC) and molecular function (MF). The color indicates the adjusted P-value, and the size indicates the gene ratio (percentage of total DEGs in each GO term). **D**. Conjoint analysis reveals a strong negative correlation between m^6^A modifications and gene expression levels (R =-0.47, p-value = 3.55e-53, Chi-squared test), with a predominant hyper-down pattern in quadrant II (64.6% of total correlations, indicating hyper-methylation paired with gene downregulation). **E**. GO term analysis of hyper-methylated and downregulated transcripts. **F**. mRNA stability assay in FXS iPSC-derived neurons and controls. RNA was isolated at designated time points following actinomycin D treatment, and the stability of STY7 mRNAs was assessed by RT-qPCR (n = 3; *** p < 0.001; two-tailed t-test).**G**. IGV tracks showing MeRIP-seq and RNA-seq read distribution for STY7 mRNAs.

### Correlation of hypermethylation with gene downregulation and increased mRNA decay

The regulatory impact of m^6^A influences RNA fate and function, modulating various aspects of RNA biology ^23–25^. However, whether aberrant m^6^A modifications are directly linked to gene expression changes in FXS neurons remains unclear. When we integrated the list of dys-methylated genes (DMGs) (Figure 5) with the DEGs (Figure 6), we revealed a strong correlation between hypermethylation and gene downregulation (“hyper-down” pattern), comprising 64.6% of total correlations (R =-0.47, p < 3.55e-53, Chi-squared test) (Figure 6D) and indicative of a mechanistic link between hypermethylation and reduced gene expression. Similarly, joint analysis of aberrantly methylated genes and DEGs from FXS postmortem tissues also demonstrated a predominant correlation between hypermethylation and gene downregulation (R =-0.33, p-value = 1.152e-53, Chi-squared test), with a notable “hyper-down” pattern in quadrant II (72.9% of total correlations) (Supplemental Figure 6J). These genes were significantly enriched in synapse-related categories (Figure 6E and Supplemental Figure 6K), suggesting that mRNA hypermethylation contributes to synaptic defects by modulating the fate of synapse-related mRNAs, in particular accelerating mRNA decay. Indeed, when we measured mRNA decay rates using actinomycin D treatment and gene-specific RT-qPCR (see Methods), we observed accelerated decay of hypermethylated and downregulated synapse-related transcripts (such as STY7) in FXS neurons (Figure 6F, G). These data suggest that hypermethylation drives transcript degradation and gene downregulation in FXS.

### m^6^A writer inhibitor rescuing synaptic and functional defects in FXS iPSC-derived cortical neurons

Given the overall increase in m^6^A levels (Figure 4A), the association of hypermethylated m^6^A marks with downregulated synapse-related genes (Figure 6D), and the observed increase in RNA decay for these hypermethylated genes (Figure 6F), we hypothesized that therapeutic intervention with an m^6^A writer inhibitor could alleviate synaptic defects in FXS neurons by restoring normal mRNA stability of synapse-related genes. To test this hypothesis, we treated FXS and control neurons with STM-2457, a small-molecule inhibitor targeting METTL3, the catalytic subunit of the METTL3/14 methyltransferase complex ^60^. MeRIP-qPCR (see Methods) revealed that excessive m^6^A marks from synapse-related transcripts in FXS neurons were reduced in the presence of STM-2457 but not DMSO (Figure 7A). Additionally, STM-2457 restored the accelerated mRNA decay of synapse-related genes such as LRRC4B in FXS iPSC-derived neurons to control levels (Figure 7B). Notably, the two-week administration of the m^6^A writer inhibitor reduced abnormal synaptic puncta and excitatory transmission, while significantly mitigating hyperexcitability in FXS neurons, without affecting normal neuronal function (Figure 7C-H). These findings highlight the critical role of m^6^A in FXS synaptic pathology and suggest therapeutic potential of m^6^A-modulating drugs in FXS.

**Figure 7.**
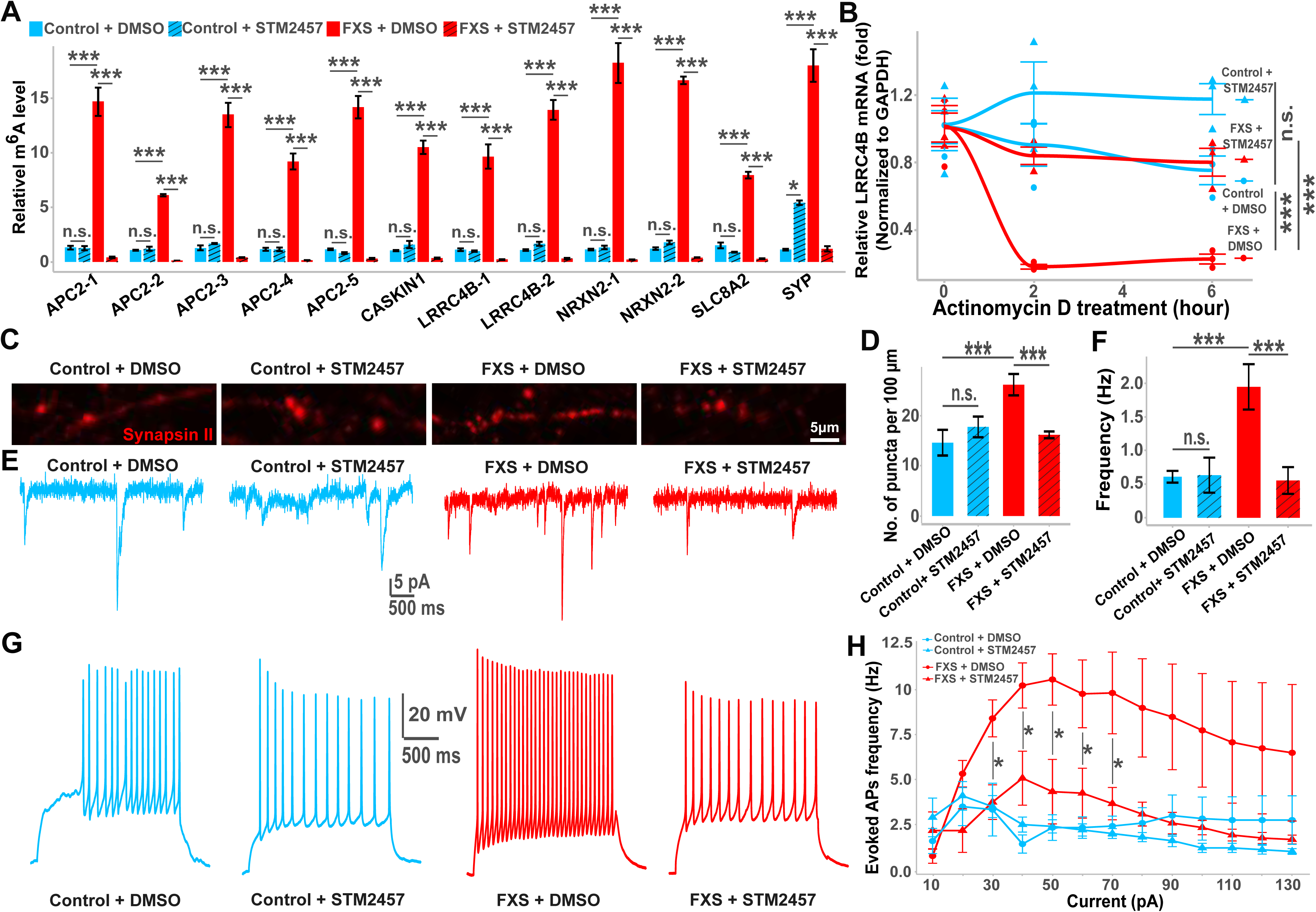
Restoration of synaptic and neuronal functions in FXS iPSC-derived cortical neurons. **A**. MeRIP-qPCR analysis showing relative enrichment of m^6^A modifications on synapse-related mRNAs in control and FXS iPSC-derived cortical neurons, with and without STM-2457 treatment. Values represent mean ± s.e.m. (n = 4 biological replicates; *p < 0.05, ***p < 0.001, n.s. denotes non-significant; one-way ANOVA with Tukey’s multiple comparisons.). **B**. mRNA stability assay in FXS iPSC-derived neurons and controls, with and without STM-2457 treatment. RNA was isolated at specified time points after actinomycin D treatment, and the stability of LRRC4B mRNAs was assessed via RT-qPCR. Values represent mean ± s.e.m. (n = 3 biological replicates; ***p < 0.001, n.s. denotes non-significant, one-way ANOVA with Tukey’s multiple comparisons.). **C-D**. Representative images and quantification of six-week-old control and FXS neurons, with and without STM-2457 treatment, stained for the synaptic marker Synapsin II. The scale bar is 5 μm. Values represent mean ± s.e.m. (n =10 to 20 neurons per condition; ***p < 0.001, n.s. denotes non-significant; one-way ANOVA with Tukey’s multiple comparisons). **E-F**. Sample whole-cell voltage-clamp recordings of excitatory postsynaptic currents (EPSCs) in six-week-old neurons, with and without STM-2457 treatment (**E**), and quantification of mean spontaneous EPSC frequencies (**F**). Values are presented as mean ± s.e.m. (n = 6 to 10 neurons per condition; ***p < 0.001, n.s. denotes non-significant; one-way ANOVA with Tukey’s multiple comparisons). **G-H**. Whole-cell current-clamp recordings of evoked action potentials (APs) (**G**), and their quantification (**H**). Values are presented as mean ± s.e.m. (n = 5 to 8 neurons per condition; *p < 0.05, n.s. denotes non-significant; two-way ANOVA with Tukey’s multiple comparisons).

## Discussion

In this report, we successfully generated iPSC lines from FXS patients matched with controls. We also demonstrated that FXS iPSC-derived neurons recapitulate key aspects of FXS pathology, including *FMR1* gene silencing, loss of FMRP, elevated synaptic protein expression, synaptic puncta, and filopodia-like spines, alongside reduced mushroom spines, consistent with previous related research ^48^. Relative to previously reported human PSCs model of FXS ^9–13^, our system is differentiated by several important features. First, beyond general synaptic transmission defects, our model identified an excitatory/inhibitory imbalance in FXS neurons.

Secondly, we characterized a disease-related hyperexcitability and hyperconnectivity in FXS cortical circuitry. Finally, and foremost, these cortical hyperexcitability and hyperconnectivity features were confirmed in the same FXS subjects via analysis of their electroencephalogram (EEG) recording. These findings confirm that cortical hyperexcitability and hyperconnectivity are key pathophysiological signatures in FXS subjects that can be modeled in patient iPSC-derived neuronal models, underscoring the translational potential and clinical relevance of our human model system in uncovering disease-related mechanisms. Taken together, our work represents the first clinically relevant human neuron model for FXS.

In the mammalian brain, m^6^A modifications are prevalent mRNA modifications^23–25^, known for their roles in brain development and neuronal function ^26–30^. Using our human FXS model, we uncovered a disease-associated increase in overall levels of m^6^A modifications and validated these findings in *Fmr1* knockout mice. Our mechanistic studies revealed that the increase in m^6^A modifications is the result of translational upregulation of m^6^A writers—including key components of the METTL3/14 complex, such as METTL3, METTL14, and WTAP—due to FMRP deficiency. These findings provide new insights into how FMRP deficiency contributes to FXS pathology. However, the selective impact of upregulated m^6^A writers on synapse-related genes in FXS requires further investigation. Additionally, whether and how FMRP deficiency directly regulates the expression of m^6^A erasers and readers awaits future mechanistic studies.

To profile transcriptome-wide m^6^A modifications, we conducted MeRIP-Seq on neurons derived from FXS patients and postmortem cortical tissues, comparing them with control samples. We identified several synapse-related genes with hypermethylation sites in both FXS neurons and postmortem tissues, suggesting that m^6^A dysregulation contributes to FXS-associated synaptic pathology. Notably, expression of many hypermethylated synapse-related genes was also downregulated, and accompanied by increased mRNA decay, indicating that m^6^A-mediated gene regulation involves modulating mRNA stability. In mice, m^6^A modifications enhances mRNA decay in neural progenitor cells to regulate normal neurodevelopment. Deficiency in m^6^A leads to premature differentiation and neurodevelopmental defects^28^. Additionally, studies in *Fmr1* knockout mice revealed that FMRP’s antagonistic interaction with the m^6^A reader YTHDF2 affects the stability of m^6^A-marked target mRNAs ^47^. Further mechanistic studies are needed to determine whether YTHDF2 reads m^6^A tags on synapse-related transcripts, accelerating their decay in FXS, and whether other m^6^A readers (e.g., YTHDF1, YTHDF3, YTHDC1, YTHDC2) modulate other mRNA functions such as alternative splicing and translational efficiency, contributing to FXS pathology.

In conclusion, our patient-derived iPSC model of FXS mimics clinically relevant cortical hyperexcitability and hyperconnectivity phenotypes. As such, our model bridges the gap between experimental models and human disease and holds paramount importance in advancing FXS research. Specifically, we uncovered a novel m^6^A-mediated epi-transcriptomic mechanism underlying FXS, where FMRP deficiency leads to increased translation of m^6^A writers, resulting in transcriptome-wide hypermethylation. This hypermethylation, in turn, disrupts the stability of synapse-related transcripts, contributing to FXS pathology. Notably, the m^6^A writer inhibitor, capable of reducing excessive m^6^A modifications, can retore normal mRNA decay and mitigate disease-related synaptic and neuronal defects in FXS iPSCs-derived neurons. Given the challenges in restoring FMRP function in FXS ^36,61^ and the notable genetic implications of the m^6^A reader in ASD ^32–35^, our findings highlight the translational potential of m^6^A modifying drug as novel therapeutic treatment for FXS and ASD. In such future directions, we envision that our FXS patient-derived iPSC model could prove useful in discovery or preclinical candidate validation stages of therapeutic development and thereby improve success rates in later-stage clinical trials.

### Limitations of the study

Several limitations remain unresolved and restrict the broader implications of our study. First, human pluripotent stem cell-derived cortical circuitry models do not fully replicate higher-order brain functions, such as cortical oscillations, due to the absence of complex brain structures and connections. This limitation prevents *in vitro* human models from fully capturing patients’ clinical phenotypes. Advanced models like human brain assembloids (e.g., cortico-striatum or cortico-thalamus assembloids) and human-mouse chimeric models (where human cortical organoids are engrafted into the cortex of immunodeficient mice) could enhance tissue organization and brain circuitry connectivity, potentially improving the modeling of higher-level brain dysfunction. Additionally, m^6^A modifications are highly dynamic and cell-specific during neurodevelopment.

Single-cell m^6^A sequencing could enable us to track these cell-specific m^6^A modifications throughout neurodevelopment, offering deeper insights into their regulatory roles in both normal cortical development and FXS pathogenesis. Lastly, only male subjects were examined in this study. Future comparisons between female and male FXS subjects could uncover sex-specific differences in m6A-mediated epi-transcriptomic dysregulation in FXS.

## Supplemental Figure legends

**Supplemental Figure 1. Modeling FXS using patient-derived iPSCs. A**. Electropherogram of Asuragen test results showing 29 CGG repeats in UHC-5 and a full mutation with over 200 CGG repeats in FXS-5 iPSC lines. **B**. Confocal images of iPSC-derived neural progenitors expressing markers such as Nestin, Sox2, ZO1, FOXG1, PAX6, and OTX2. The scale bar is 50 μm. **C**. Quantification of neural progenitor marker expression. Values represent mean ± s.e.m. (n = 5 to 76 neurons per condition). **D-L**. Semi-automated Sholl analysis reveals no significant differences in neurite parameters between FXS patient-derived cortical neurons and control neurons. Shown are representative images of GFP-labeled FXS and control neurons (**D**), a schematic of neurite analysis (**E**), and quantification of neurite parameters including number of sections (**F**), total neurite length (**G**), terminal points (**H**), branch points (**I**), maximum section length (**J**), minimum section length (**K**), and mean section length (**L**). The scale bar is 50 μm. Values represent mean ± s.e.m. (n = 30 to 169 neurons per condition; n.s. denotes non-significance; two-tailed t-test).

**Supplemental Figure 2. Sodium and potassium channel activity. A-B**. Quantification of evoked action potential (AP) amplitude (**A**) and rheobase (**B**). Values are presented as mean ± s.e.m. (n = 43 to 92 neurons per condition; n.s. denotes non-significance; two-tailed t-test). **C-E**. Sample whole-cell voltage-clamp recording traces of sodium and potassium channel activity (**C**), along with quantification of sodium (**D**) and potassium (**E**) currents. Values are presented as mean ± s.e.m. (n = 56 to 107 neurons per condition; n.s. denotes non-significance; two-tailed t-test).

**Supplemental Figure 3. MEA and EEG recordings. A**. Sample bright-field images of human cortical neuron cultures at 6 weeks post-neuronal differentiation. The scale bar is 50 μm. **B-C**. Increased waveforms detected per electrode by automatic spiking sorting in FXS cortical microcircuitry compared to controls. Shown are sample waveforms for electrode 6 and 26 in control and FXS iPSCs-derived cortical microcircuitry (**B**). Quantification of waveforms per channel is also presented (**C**). Values represent mean ± s.e.m. (n = 3 cultures; ***p < 0.001; two-tailed t-test). **D**. Power spectral density (PSD) traces across different frequency bands from four typically developing control (TDC) and FXS subjects’ EEG data. Numbers indicate TDC and FXS experimental IDs.

**Supplemental Figure 4. Elevated m^6^A levels associated with increased m^6^A writer expression in *Fmr1* knockout mice. A**. Increased m^6^A levels in *Fmr1* knockout mice compared to wild type (WT) mice. Shown are dot blot assays of polyA^+^ RNA m^6^A levels and quantification of m^6^A intensity in *Fmr1* knockout neurons relative to WT, normalized to methylene blue (n = 4 samples; ***p < 0.001; two-tailed t-test). **B**. Transcriptional expression analysis revealing no significant change of METTL3, METTL14, and WTAP transcripts from RNA sequencing between WT and *Fmr1* knockout mice. Values represent mean ± s.e.m. (n = 2 wild type and 2 *Fmr1* knockout mice; n.s. denotes non-significance; two-tailed t-test). **C-D**. Western blot analysis and quantification of FMRP, METTL3, and WTAP protein levels. Values represent mean ± s.e.m. (n = 4 samples; ***p < 0.001; two-tailed t-test).

**Supplemental Figure 5. Transcriptome-wide m^6^A dysregulation in postmortem brain tissues from FXS subjects. A-C**. Transcriptome-wide m^6^A profiling in postmortem brain tissue (Brodmann area 44) derived from FXS subjects and healthy controls. **A**. Metagene plot showing m^6^A peak distribution across human chromosomes. **B**. Sequence logo depicting the top enriched motif in m^6^A peaks (p value = 10⁻⁴). **C**. Distribution of m^6^A methylation across transcripts, with each transcript divided into 5′ UTR, CDS, and 3′ UTR. **D**. Volcano plot displaying MeRIP-seq data of differentially methylated m^6^A peaks between FXS and control groups (n = 3 healthy controls and 3 FXS samples; adjusted p-value < 0.01; blue and red dots indicate significant hypo-and hyper-methylated sites, respectively). **E-F**. Histogram illustrating the distribution of hypo-and hyper-methylated sites per gene (**E**) and per chromosome length (**F**), with blue and red boxes indicating hypo-and hyper-methylated sites, respectively. **G**. Gene Ontology (GO) term analysis of differentially methylated genes (DMGs) between FXS subjects and controls, showing top 15 enrichment terms for cellular component (CC) and biological process (BP). The color represents the adjusted P-value, and the size of the shape represents the gene ratio (percentage of total DMGs in the given GO term). **H-J**. Venn diagrams showing the overlap between hypermethylated transcripts and FMRP target transcripts (**H**), SFARI ASD genes (**I**), and reported synapse-related genes (**J**), along with representation factors and p-values. **K-L**. GO term analysis of overlapped genes between hypermethylated transcripts and FMRP target transcripts (**K**) and between hypermethylated transcripts and SFARI ASD genes (**L**).

**Supplemental Figure 6. Correlation of hypermethylation with gene downregulation in postmortem brain tissues from FXS subjects. A**. Venn diagram, drawn to scale, shows the overlap between differentially expressed genes (DEGs) in FXS iPSCs-derived neurons and controls, and FMRP target transcripts, with representation factor and P-value. **B**. Gene Ontology (GO) term analysis of overlapped genes between hypermethylated transcripts and FMRP target transcripts reveals the top 15 enriched terms for cellular component (CC) and biological process (BP). The color represents the adjusted p-value, and the size represents the gene ratio, indicating the percentage of total DMGs within each GO term. **C**. Venn diagram shows the overlap between DEGs and reported synaptic genes, with representation factor and P-value. **D**. Volcano plot illustrating differential gene expression analysis in postmortem brain tissue (Brodmann area 44) from FXS subjects versus healthy controls (n = 3 healthy controls and 3 FXS samples; fold change > 1.4, p-value < 0.05; blue and red dots represent significantly down-and up-regulated genes, respectively). **E-F**. Venn diagrams show the overlap between DEGs from postmortem tissues and FMRP target transcripts (**E**) and SFARI ASD genes (**F**), along with representation factors and P-values. **G-H**. GO term analysis of overlapped genes between DEGs and FMRP target transcripts (**G**), as well as DEGs and SFARI ASD genes (**H**), revealing the top 15 enriched terms for cellular component (CC), biological process (BP), and molecular function (MF). **I**. Venn diagram illustrating the overlap between DEGs from postmortem tissues and reported synaptic genes, with the representation factor and P-value. **J**. Conjoint analysis shows a strong negative correlation between m^6^A modifications and gene expression levels from postmortem brain tissues (R =-0.33, p-value = 1.152e-53, Chi-squared test), with a notable hyper-down pattern in quadrant II (hyper-methylation paired with gene downregulation, 72.9% of total correlations). **K**. GO term analysis of transcripts with hyper-methylation paired with gene downregulation.

## Acknowledgements

We would like to thank to the members of Guo Laboratory, Dr. Chuan He, as well as Drs. Aaron Zorn, James Well, and Kenneth Campbell, for their valuable comments and suggestions. We also thank Dr. Andrew Volk for providing the WTAP antibody. This study has received support from the National Institutes of Health (NIH) grants (R21NS122169 and R21MH132038), the Cincinnati Children’s Hospital Medical Center (CCHMC) Trustee Grant Award, and the Local Initiative for Excellence Foundation to Dr. Ziyuan Guo. The NIH grant R01NS115507 and Department of Veteran Affairs grants I01BX002149 and IK6BX006316 to Dr. Haining Zhu are acknowledged. We would also like to express our gratitude to NIH NeuroBioBank for providing human postmortem brain tissues.

## Author Contributions

Author Contributions: L.L. led and was involved in every aspect of the project. A.S. conducted whole-cell patch-clamp recording and analysis. L.D., X.C., and J.T. contributed to cell cultures and immunostaining. J.L., M.M., and E.P. conducted EEG analysis. C.M., P.T., and H.Z. performed polysome profiling analysis. G.Y., A.L., and C.W made contributions to data analysis. C.W., L.S., and C.E. provided clinical samples and data. M.C. and M.S. carried out Asuragen testing. Y.Z. contributed to MEA analysis. Y.L. and N.S. made contributions to MeRIP sequencing and RNA sequencing analysis. X.W., L.Z., and D.L. contributed to experimental design. C.G. made contributions to immunoprecipitation experiments and provide *Fmr1* knockout brains. Z.G. conceptualized and designed the project and wrote the manuscript.

## Author information

MeRIP sequencing and RNA sequencing data were deposit at GEO. Correspondence and requests for materials should be address to Z.G.

## Methods

### Participants

Seven male participants with a full mutation in the *FMR1* gene were recruited through the Cincinnati Fragile X Research and Treatment Center. Fragile X status was confirmed by southern blotting and PCR analysis completed at Rush University. Five male typically developing control (TDC) were recruited through web-based fliers from the local community and had no prior diagnosis or treatment for developmental or neuropsychiatric disorders. Written informed consent was obtained from all participants or their legal guardians, with verbal assent provided when appropriate. This study was approved by the Institutional Review Board (IRB) of Cincinnati Children’s Hospital Medical Center (CCHMC) and was conducted in accordance with all relevant guidelines and regulations, including the Declaration of Helsinki.

Deviation intelligence quotient (IQ) scores, including full-scale IQ (FSIQ), non-verbal IQ (NVIQ), and verbal IQ (VIQ), were assessed using the Stanford-Binet Intelligence Scales, 5th Edition, with adjustments made to account for floor effects commonly seen in individuals with developmental disabilities ^1^.

Whole blood samples were collected from participants for various analyses: one set was used for peripheral blood mononuclear cells (PBMCs) isolation and FXS induced pluripotent stem cells (iPSCs) generation which will be detailed in the subsequent section, another for FMRP quantification using an optimized Luminex assay as described by Boggs *et al.* ^2^, and a third for *FMR1* mRNA detection via a sensitive RT-qPCR assay as previously described ^3^.

### Generation and culture of iPSC lines

PBMC were isolated from one set of whole blood samples using SepMate^TM^-50 tubes (StemCell Technologies) with Lymphoprep^TM^ Density Gradient Medium (StemCell Technologies) through density-gradient centrifugation. A total of 500,000 PBMCs were seeded in 24-well plates and cultured in PBMC medium [StemPro^®^-34 SFM medium (ThermoFisher) supplemented with SCF (100ng/mL; Peprotech), FLT-3 (100ng/mL; Peprotech), IL-3 (20ng/mL; Peprotech), and IL-6 (20ng/mL; Peprotech)] for four days. PBMCs were then transduced using the Cytotune™-iPS 2.0 Sendai Reprogramming Kit (ThermoFisher) at the appropriate MOI, following the manufacturer’s instructions, and incubated overnight. The next day, the CytoTune™ 2.0 Sendai viruses were removed, and the transduced PBMCs were cultured in PBMC medium for two additional days. The cells were subsequently seeded on vitronectin (ThermoFisher)-coated plates in complete StemPro^®^-34 SFM medium without cytokines for four days. After this, the medium was switched to Essential 8™ Flex medium (ThermoFisher). Once iPSC-like colonies emerged, they were manually picked and cultured in Essential 8™ Flex medium. These FXS iPSCs were labeled FXS-1, FXS-2, FXS-3, FXS-4, FXS-5, FXS-6, FXS-7, separately. The control iPSCs were named UHC-1, UHC-2, UHC-3, UHC-5, UHC-4, UHC-6, separately.

All iPSCs were expanded on vitronectin-coated plates in Essential 8™ Flex medium and passaged with ReleSR^TM^ (StemCell Technologies) upon reaching 70-80% confluence using a 1:6 split ratio. iPSCs were cryopreserved in stem cell freezing medium [American Type Culture Collection (ATCC)]. 10 µM Y27632 (Peprotech) was added during both passaging and thawing. The karyotype of all iPSCs generated in this project have been characterized by the Pluripotent Stem Cell Facility at CCHMC. And their expression of stemness markers have been confirmed by immunostaining.

### CGG sizing

Genomic DNA was extracted from iPSCs using the Quick-DNA Miniprep Kit (Zymo Research). The CGG repeat size in the *FMR1* gene was determined by repeat-primed PCR (RP-PCR) using primers flanking the CGG-repeat region of the 5’ UTR in *FMR1*, as per the manufacturer’s instructions (AmplideX ^®^ PCR/CE *FMR1* Kit, Asuragen). Briefly, 2 μl of purified genomic DNA (40 ng/µl) was added to a master mix containing 11.45 µl GC-Rich Amp Buffer, 0.5 µl *FMR1* F, R FAM-Primers, 0.5 µl *FMR1* CGG Primer, 0.5 µl diluent, and 0.05 µl GC-Rich Polymerase Mix. The PCR protocol included: initial denaturation at 95 °C for 5 min, followed by 10 cycles of 97 °C for 35 s, 62 °C for 35 s, and 68 °C for 4 min. This was followed by 20 cycles of 97°C for 35 seconds, 62°C for 35 seconds, and 68°C for 4 minutes, with an additional 20-second increment per cycle. The final extension was carried out at 72°C for 10 minutes. The PCR products were then mixed with Hi-Di Formamide and the ROX 1000 Size Ladder. After denaturation at 95 °C for 2 min and cooling to 4°C for 5 minutes, the amplicons were separated via capillary electrophoresis (CE) using an ABI 3500 Genetic Analyzer with POP-7 polymer (Applied Biosystems). The CGG repeat length was calculated based on the fragment size in base pairs using the *FMR1* Analysis Macro (v2.1.2; Asuragen) with size and mobility conversion factors.

### Differentiation of cortical neurons from iPSCs

Cortical neurons were generated from iPSCs as previously described^6, 7^, with some modifications as described here. Briefly, iPSCs were dissociated into single cells with StemPro™ Accutase™ Cell Dissociation Reagent (ThermoFisher) and seeded at a density of 2,500,000 cells per well on Matrigel-coated (human embryonic stem cell-qualified matrix, Corning) six-well plates in Essential 8™ Flex medium with Y27632. From day 1 to day 3, the medium was switched to Essential 6™ medium (ThermoFisher) supplemented with LDN193189 (500 nM, Peprotech), SB431542 (10 µM, Cayman Chemical), and XAV939 (5 µM, Selleckchem) (referred to as E6-LSBX medium). From day 4 to day 8, cells were maintained in E6-LSB medium (without XAV939). Beginning on day 9, the medium was switched to a house-made N2 medium [DMEM/F12 medium with 2-mercaptoethanol (1X; ThermoFisher), sodium bicarbonate (2 mg/mL; Sigma), D-(+)-Glucose (1.56 mg/mL; Sigma), progesterone (6.4 mg/mL; Sigma), N2 Supplement-B (0.5X; StemCell Technologies)], supplemented with B27™ (1X; ThermoFisher) and FGF-8b (50ng/mL; R&D Systems) until day 21. At this stage, neuronal progenitor cells (NPCs) were either cryopreserved in stem cell freezing medium or dissociated with Accutase™ and plated onto Poly-L-ornithine (PO, 15 µg/mL; Sigma)/Laminin (2 µg/mL; R&D Systems)/Fibronectin (1 µg/mL; R&D Systems)-coated plates or coverslips. For quantifying numbers of cells expressing neural progenitor cells and cortical neuronal markers, 50,000 cells were seeded onto coverslips. For protein and RNA extractions, 5,000,000 cells were seeded onto 10 cm dishes. Cells were then cultured in neuron medium [Neurobasal® medium (ThermoFisher) with B27™ supplement (1X), BDNF (20 ng/mL; Peprotech), GDNF (20 ng/mL; Peprotech), Ascorbic Acid (AA, 0.2 mM; Sigma), cAMP (100 µM; Sigma), GlutaMAX™ (1X; ThermoFisher), and MEM Non-Essential Amino Acids (NEAA, 1X; ThermoFisher)] with the addition of DAPT (10 µM; Cayman Chemical). Half of the neuron medium was replaced twice a week for a total of six weeks. For treatment, cultures were exposed to either DMSO (0.1%; Sigma) or 10 μM STM2457 (Cayman Chemical) for 2 weeks, from week 4 to week 6.

### Isolation of primary mouse astrocytes and co-culture with iPSC-derived cortical neurons

Primary mouse astrocytes were isolated as previously described ^8^, with minor modifications. Brains from P0 C57BL/6J mice were dissected, and the meninges were removed. The meninges-free cerebral cortices were minced into small pieces and digested with trypsin (ThermoFisher) and DNase I (Sigma). The resulting cell suspension was plated onto 0.01% Poly-L-Lysine (PLL; Sigma)-coated flasks in Dulbecco’s Modified Eagle Medium (DMEM, ThermoFisher) supplemented with 10% fetal bovine serum (FBS) (ThermoFisher). and 2% penicillin/streptomycin. From day 2 to day 8, primary astrocytes were separated from oligodendrocyte progenitor cells, microglia and neurons by tapping the flask and washing with PBS. Primary astrocytes were cryopreserved around day 8.

Before co-culturing with cryopreserved iPSC-derived NPCs, primary mouse astrocytes were thawed, expanded and seeded on coverslips in 24-well plates. Once the astrocyte monolayer reached 80-90% confluence, cryopreserved iPSC-derived NPCs were thawed, 20,000 cells were seeded directly onto the astrocytes, and differentiated in neuron medium [Neurobasal® medium supplemented with B27™, BDNF, GDNF, AA, cAMP, GlutaMAX™, and NEAA] containing DAPT. Half of the neuron medium was replaced twice weekly. After 6 weeks of culture, whole-cell patch clamp recordings and immunocytochemistry for synaptic puncta and neurites were performed. For treatment, cultures were exposed to either DMSO (0.1%; Sigma) or 10 μM STM2457 (Cayman Chemical) for 2 weeks, from week 4 to week 6.

### Western blot analysis

Ten-week-old wild-type (WT) and *Fmr1*-knockout (*Fmr1*-KO) male mice, kindly provided by Prof. Christina Gross, were used for protein extraction from fresh cortical tissues. Additionally, protein was extracted from six-week-old iPSC-derived cortical neurons using ice-cold RIPA buffer (Sigma), supplemented with cOmplete Protease Inhibitor Cocktail (Roche) and PhosSTOP^™^ (Roche). Lysates were sonicated and incubated on ice, followed by centrifugation at 4°C. Protein concentrations were determined using the Pierce™ BCA Protein Assay Kit (ThermoFisher). Equal amounts of protein lysates were denatured in SDS-polyacrylamide gel electrophoresis (PAGE) Sample Loading Buffer (G-Biosciences), separated on 4-12% Bolt Bis-Tris Plus Protein Gels (ThermoFisher), and transferred to polyvinylidene fluoride (PVDF) membranes (Millipore) using the XCell SureLock™ Mini-Cell system (ThermoFisher). The membranes were blocked for 1 hour at room temperature (RT) with 5% non-fat milk in Tris-buffered saline containing Tween-20 (TBST) (blocking buffer), followed by overnight incubation at 4°C with the following primary antibodies diluted in blocking buffer: rabbit polyclonal anti-FMRP (1:1000; Abcam), rabbit monoclonal Synapsin II (1:1000; Abcam), mouse monoclonal SV2 (1:1000; DSHB), rabbit monoclonal METTL3 (1:500; MedChemExpress), rabbit polyclonal METTL14 (1:1000; Sigma), rabbit monoclonal WTAP (1:500; Cell Signaling Technology), and mouse monoclonal GAPDH (1:5000; Santa Cruz Biotechnology). After 3 washes with TBST, the membranes were incubated for 1 hour at RT with horseradish peroxidase (HRP)-coupled secondary antibodies: anti-rabbit IgG (1:5000; ThermoFisher) or anti-mouse IgG (1:5000; ThermoFisher). The membranes were washed 3 times with TBST and visualized by the addition of SuperSignal™ West Pico PLUS Chemiluminescent Substrate (ThermoFisher) directly on the membranes placed in the iBright^™^ FL 1500 Imaging System (ThermoFisher). Protein levels were normalized to GAPDH, and quantitative analysis was performed using ImageJ software.

### Immunocytochemistry

Six-week-old iPSC-derived cortical neurons were fixed with 4% paraformaldehyde (ThermoFisher) for 15 min at RT, then washed 3 times with PBS. They were permeabilized with 0.1% Triton^™^ X-100 (Sigma) and blocked in 3% donkey serum (Sigma) for 2 hours. Primary antibodies, diluted in PBS containing 0.1% Triton X-100 and 3% donkey serum, were applied to the cells and incubated overnight at 4°C. Following primary antibody incubation, cells were treated with species-specific secondary antibodies (1:1000; Jackson ImmunoResearch Laboratories) diluted in 3% donkey serum for 2 hours at RT. Cells were washed and cell nuclei were stained with 4′,6-diamidino-2-phenylindole (DAPI) for 5 min. Coverslips were mounted in fluoroshield mounting medium (Abcam), sealed with clear nail polish, and stored at 2-8°C until imaging. The following primary antibodies were used: mouse monoclonal Nestin (1:500, Santa Cruz Biotechnology), mouse monoclonal Sox2 (1:1000; Santa Cruz Biotechnology), mouse monoclonal ZO1 (1:1000; Thermo Fisher Scientific), rabbit polyclonal Foxg1 (1:1000, Abcam), mouse monoclonal PAX6 (1:1000; BD Biosciences), mouse monoclonal OTX2 (1:50; DSHB), rabbit polyclonal doublecortin (1:1000; Cell Signaling Technology), mouse monoclonal BRN2 (1:50; DSHB), goat monoclonal CaMKII (1:500; Santa Cruz Biotechnology), mouse polyclonal beta III tubulin (1:500; Sigma), rabbit polyclonal TBR1 (1:1000; Abcam), rat monoclonal CTIP2 (1:1000; Abcam), rabbit monoclonal Synapsin I (1:1000; Millipore), mouse monoclonal SV2 (1:1000; Santa Cruz Biotechnology).

Images were obtained on either a Nikon A1R inverted confocal laser scanning microscope with 60x objectives or an Olympus BX51 with 20x objectives. Image analysis was performed using NIS Elements version 5.42.06 software (Nikon) and Imaris version 10.2.0 software (Nikon). For quantifying NPC and neuron markers, the “spot” function was used. For neurite analysis, the neurite structures were identified using the “filament” function, parameters including the number of segments, total length, terminal points, branch points, maximum segment length, minimum segment length, and mean segment length were assessed. Synaptic puncta were identified using the “spot” function, and only the puncta located around the dendritic and axonal filaments were selected for quantification. Synaptic density (D) was determined by calculating the total SV2 and SYN1 puncta per 100 µm of total dendritic length.

### Whole-cell patch clamp recording

Whole-cell patch-clamp recordings were conducted at RT on iPSC-derived cortical neurons, either untreated or treated with DMSO or STM2457 for 2 weeks, following a six-week co-culture with primary mouse astrocytes, using a Multiclamp 700A patch-clamp amplifier (Molecular Devices) as previously described ^9^. Briefly, cortical neurons grown on coverslips were visualized with a 40X air objective on an inverted Zeiss microscope, and the recording chamber was constantly perfused with a bath solution consisting of 128 mM NaCl, 30 mM glucose, 25 mM HEPES, 5 mM KCl, 2 mM CaCl_2_, and 1 mM MgCl_2_ (pH 7.3; 315–325 Osm). Patch pipettes were pulled from borosilicate glass (Sutter Instrument) with a resistance of 3-5 MΩ and filled with an internal solution composed of 135 mM CsGluconate, 10 mM Trisphosphocreatine, 10 mM HEPES, 2.5 mM EGTA, 4 mM MgATP, 0.5 mM Na₂GTP, and 5 mM QX-314 (pH 7.3 with CsOH; ∼305 mOsm) for excitatory postsynaptic currents (EPSCs). EPSCs were recorded at a holding potential of-70 mV. The series resistance was typically between 10-30 MΩ. For spontaneous EPSC recordings, 20 µM bicuculline (Tocris Bioscience) was added. Drug application was achieved via a gravity-driven drug delivery system (VC-6, Warner Instruments). Spontaneous EPSCs were recorded for 4 minutes (starting 2 minutes after break-in, to allow QX-314 to block sodium currents) at a holding potential, sampled at 10 kHz, and filtered at 1 kHz. Spontaneous and evoked action potentials (AP) were recorded using current clamp, while sodium and potassium currents were measured using voltage clamp. Data acquisition was performed using pClamp 9 software (Molecular Devices), and spontaneous synaptic events were analyzed with MiniAnalysis software (Synaptosoft).

### Microelectrode array (MEA) and analysis

The 64-channel multielectrode array (MEA) recording system (MED64, Alpha MED Scientific) features an array of 64 planar microelectrodes arranged in an 8 × 8 pattern (50 × 50 μm electrodes with 150 μm spacing) and was used to non-invasively monitor the electrical activity of neuronal networks. Before use, the surface of the MED64 probe was coated with PO/laminin/fibronectin and a drop of iPSC-derived NPCs (500,000 cells) were directly plated on the top of the electrodes and cultured in B-27™ Electrophysiology Kit supplemented with BDNF (20 ng/ml), GDNF (20 ng/ml), AA (0.2 mM), cAMP (100 µM), GlutaMAX (1X), and NEAA (1X).

DAPT was administered on day 1 to synchronize neuronal maturation, and the cultures were fed weekly. Spontaneous neuronal activity was recorded at week 8post-differentiation for 5 minutes at a sampling rate of 20 kHz using Mobius software (Alpha MED Scientific). Data were filtered with a 1-1000 Hz frequency bandwidth.

A spike was detected when the recorded signal exceeded a threshold of ± 5.5 σ, where σ was the standard deviation (SD) of the baseline noise during quiescent periods. Spike clustering was performed using the Mobius software’s ‘Cluster Spikes’ panel to distinguish and separate signals from multiple cells recorded on a single channel based on the shapes of their spikes, utilizing a modified version of the online ‘Leader-Follower Clustering’ algorithm. A valid cluster was defined by the following parameters: a minimum of 5 spikes (‘Min # spikes’) and 30% similarity (‘Similarity %’). Next, burst activity and synchronized burst activity were analyzed by Mobius offline toolkit software (Alpha Med Scientific). Bursts were identified as sequences of at least three spikes with inter-spike intervals within 300 milliseconds and a duration exceeding 200 milliseconds. Burst duration and spikes per burst were analyzed and compared.

Synchronized burst activity, representing burst events across 64 channels, was visualized using a ‘time course of spike frequencies across 64 channels’ plot, where spike frequencies were quantified as the array-wide spike detection rate (ASDR). Additionally, a raster plot was used to display the distribution of spike timing across all channels.

Moreover, MEA data were analyzed using UP state analysis, a method previously applied to *in vitro* brain slices from *Fmr1*-KO mice ^10^. Spontaneous UP states were detected based on methods defined in Hays et al^10^. For each channel, data were normalized to zero by subtracting average, and rectified. Rectified data were then low pass filtered at 0.2 Hz. UP states were identified when the amplitude of the filtered signal exceeded the root mean square threshold for a minimum of 200 milliseconds. Events within 600 milliseconds of each other were grouped together into a single UP state. Median duration of up states as well as interval between up states was calculated for each channel.

### EEG recording, data preprocessing, and analysis

Five minutes of resting state EEG data was collected from each participant. To facilitate cooperation in FXS patients, participants watched a silent video during recording. EEG data were recorded with a 128-electrode saline-based Hydrocel system (Magstim/EGI) at a sampling rate of 1000 Hz.

EEG data were blinded, and all preprocessing was performed using EEGLAB (EEGLAB)^11^. The data were filtered offline using a bandpass filter from 0.5 to 120 Hz, and a notch filter at 60 Hz was used to remove power line noise. EEG data was visually inspected for channels or periods affected by excessive artifacts from muscle, movement, and channel noise. Removed channels were interpolated using spherical spline interpolation. Independent component analysis (ICA) decomposition was performed using the EEGLAB pop_runica function, and components were visually inspected to identify and remove those containing muscle, channel, and eye movements. The data were then referenced to average.

Preprocessed EEG data were analyzed using MNE (v 1.6.1) and the FOOOF toolbox (v 1.1.0). Power spectral density (PSD) was calculated at each electrode using Welch’s method, with two second segments and 50% overlap. For each subject, the PSD was averaged across all channels excluding two outer rows of peripheral channels, to minimize the effect of remaining muscle and movement artifact. The FOOOF toolbox was used to model the periodic and aperiodic components of the average PSD for each subject. The model was computed using the fixed aperiodic mode, with peak width limits of 1 to 12 Hz, maximum peak number set to 6, peak threshold of 2 standard deviations, and no minimum peak height.

### m^6^A dot blot assay on cortical neurons and *Fmr1*-KO mouse brain tissue

Ten-week-old WT and *Fmr1*-KO male mice were used for RNA extraction from fresh cortical tissues. Additionally, total RNA was extracted from six-week-old iPSC-derived cortical neurons using TRIzol reagent (ThermoFisher), followed by purification with Dynabeads mRNA Purification Kit (ThermoFisher) according to the manufacturer’s instructions. The extracted mRNA was denatured and spotted onto Hybond N+ membrane (GE Healthcare), then cross-linked using a Stratalinker UV Crosslinker 2400 (Stratagene). The membrane was blocked with 5% milk in PBST and probed overnight at 4°C with a rabbit m^6^A antibody (1:250; Synaptic Systems), followed by incubation with HRP-conjugated anti-rabbit IgG secondary antibody (1:5000). Detection was carried out using SuperSignal™ West Pico PLUS Chemiluminescent Substrate, and images were captured using the iBright™ FL 1500 Imaging System. To determine the total RNA input, the membrane was stained with 0.04% methylene blue (Sigma) in 0.5M sodium acetate (Millipore) prior to blocking. Quantification of dot intensity was performed using ImageJ software.

### MeRIP-seq, MeRIP-RT-qPCR and analysis

The mRNAs from total RNAs were isolated from six-week-old cortical neurons using Dynabeads mRNA Purification Kit. Purified mRNA samples were fragmented to 100-200 nt with 10X RNA Fragmentation Buffer (100 mM Tris-HCl, 100 mM ZnCl_2_ in nuclease-free H_2_O). The reaction was stopped by adding 10X EDTA (0.5M). 10% of the fragmented RNA was set aside as an input sample, while the remaining RNA was used for m^6^A immunoprecipitation. The fragmented RNA was incubated with anti-m^6^A antibody (Synaptic Systems) for 3 h at 4°C and then with protein G magnetic beads (ThermoFisher) at 4°C for an additional 2 h to obtain immunoprecipitated RNA fragments. The m^6^A-enriched RNA was extracted using TRIzol. The library was prepared following the instructions for Kapa RNA HyperPrep Kit (Roche). Both the input samples without IP and the m^6^A IP samples were subjected to 150-bp, paired-end sequencing on an Illumina NovaSeq 6000 sequencer at CD Genomics.

For MeRIP-seq analysis, quality control of raw sequencing reads was assessed using FastQC (v0.11.7). Raw reads in fastq format were then subjected to trimming, eliminating adaptor sequences, reads with adaptor contaminants, and low-quality reads, using Trim Galore (v0.6.6). The preprocessed reads were aligned to the human genome (GRCh38) employing STAR (v2.7.9) with default settings. m^6^A peaks were identified using the R package exomePeak (v2.16.0) with the following parameters: “PEAK_CUTOFF_PVALUE = NA, PEAK_CUTOFF_FDR = 0.05, FRAGMENT_LENGTH = 200”. The distributions of m^6^A peaks across the chromosomes in both UHC and FXS cortical neurons were analyzed using circos (v0.69.8). Differential m^6^A peak were identified using exomePeak R package under parameters: “PEAK_CUTOFF_PVALUE = NA, PEAK_CUTOFF_FDR = 0.05, FRAGMENT_LENGTH=200”.

The output “sig_diff_peak” file was further filtered using the condition “|diff.lg.fdr| > 2” and “|diff.log2.fc| > 0”, generating a highly significant differential m^6^A peak file between FXS and UHC neurons (Table S2). This dataset was then categorized into hypermethylated and hypomethylated m^6^A peaks based on the diff.log2.fc values. Peaks with “diff.log2.fc” > 0 were classified as hypermethylated, while those with “diff.log2.fc” < 0 were classified as hypomethylated (Table S2 and S4). Identified m^6^A peaks were visualized using the Integrative Genomics Viewer (IGV, v2.15.4) with bam formats. Gene ontology (GO) analysis was performed using the clusterProfiler R package (v4.11.0), with visualizations generated in RStudio (v3.4.3) using the ggplot2 package (v3.5.1). In the plots, circle size corresponds to the percentage of genes in each ontology group, and color represents the p-value for gene enrichment associated with each GO term. m^6^A-RNA-related genomic features were visualized using Guitar R package (v2.14.0), and de novo motif analysis was performed with homer (v4.11).

To examine m^6^A modifications on individual genes in six-week-old cortical neurons, MeRIP-RT-qPCR was performed using the same procedures, except that RNAs were sheared into approximately 200 nt fragments. Total RNAs were isolated from neurons untreated or treated with DMSO or STM2457 for 48 hours. Primers were designed to target different m^6^A modification sites identified through MeRIP-seq data, including 5 sites for *APC2*, 1 site for *CASKIN1,* 2 sites for *LRRC4B,* 2 sites for *NRXN2,* 1 site for *SLC8A2,* 1 site for *SYP,* 1 site for *SYT3*, and 1 site for *SYT5.* All primers used for MeRIP-RT-qPCR are listed in Table S1. RT-qPCR was conducted using PowerUp SYBR Master Mix on a QuantStudio system. m^6^A enrichment was determined by RT-qPCR analysis, calculating the percentage of a target gene in IP fraction relative to the input fraction using the formula: %Input = 2^˄^ (Ct of target gene in IP - adjusted Ct of target gene in input).

### RIP-RT-qPCR

HEK293T cells were grown in DMEM supplemented with 10% FBS and transfected using Lipofectamine 3000 (Thermo Fisher Scientific) according to the manufacturer’s protocol. Cells were either co-transfected with *FMRP-Flag* and *FUGW* plasmids or transfected with *mCherry-Flag* plasmids. Post-transfection, cells were lysed using 1 mL of RIP lysis buffer (10 mM HEPES pH 7.4, 200 mM NaCl, 30 mM EDTA, 0.5% Triton X-100) supplemented with protease inhibitors and SUPERaseIN (1:500; ThermoFisher) at 4°C with rotation for 30 minutes. Cell lysates were then cleared by centrifugation at 20,000× g at 4°C for 15 min. 10% of the supernatant was saved for RNA extraction and protein extraction as input controls. The remaining supernatants were incubated overnight at 4°C with ANTI-FLAG^®^ M2 Affinity Gel (Sigma) with rotation. The immunoprecipitated protein-RNA complexes were subsequently washed three times with RIP wash buffer (10 mM HEPES, pH 7.4, 400 mM NaCl, 30 mM EDTA, 0.5% Triton X-100, 2X protease inhibitors, and 1:1000 SUPERaseIN). Saved input and immunoprecipitated proteins were eluted with sample loading buffer for SDS-PAGE, followed by western blotting as described above using rabbit polyclonal FMRP (1:1000; Abcam) and mouse monoclonal GAPDH (1:5000; Santa Cruz Biotechnology). The saved input and immunoprecipitated RNA were extracted with TRIzol reagent (ThermoFisher) and subjected to RT-qPCR with primers (listed in Table S1), normalizing to input RNA. Reverse transcription was performed using the High-Capacity cDNA Reverse Transcription Kit (ThermoFisher), and RT-qPCR was conducted using PowerUp SYBR Master Mix (ThermoFisher) on a QuantStudio system.

### Polysome profiling and polysome-associated mRNA analysis

Sucrose density gradient centrifugation was employed to separate the ribosome fractions as described previously ^12^. Briefly, six-week-old cortical neurons were cultured with 100 μg/mL cycloheximide (CHX; Sigma) supplement for 10 min at 37 °C prior to cell harvest. Cells were washed in ice-cold PBS containing 100 μg/mL CHX and harvested in polysome lysis buffer (5 mM Tris-HCl, pH7.5, 2.5 mM MgCl2, 1.5 mM KCl, 1 mM DTT,0.5% Triton X-100, 0.5% sodium deoxycholate, 100 μg/mL CHX, 40 U/ml RNaseOUT, 0.2 mg/mL heparin and protease inhibitor cocktail). Cells were lysed by pipetting several times and incubation on ice for 15 min, followed by centrifugation at 10,000 x g for 10 min at 4 °C. 10–50% linear sucrose gradient was prepared using Biocomp Gradient Master 108 with pre-layered 10% and 50% sucrose in the gradient buffer (5 mM Tris-HCl, pH7.5, 2.5 mM MgCl2,1.5 mM KCl, 2 mM DTT, 100 μg/mL CHX, 40 U/ml RNaseOUT, 0.1 mg/mL heparin and protease inhibitor cocktail). 300 μl supernatant of cell lysate was layered on the top the sucrose gradient and centrifuged at Beckman SW41 rotor in Beckman Ultra-clear tube at 4 °C, 39K rpm for 2 hrs. Gradients were fractionated while monitoring absorbance at A260 with a Biocomp Piston Gradient Fractionator system according to the manufacturer protocol. The areas of monosome fractions (#6-11) and heavy polysome fractions (#18-25) were calculated using GraphPad Prism 6 (GraphPad Software Inc.). Total RNA in each fraction was isolated using TRIzol Reagent. 200 ng of RNA in each fraction was reverse transcribed to cDNA using a High-Capacity cDNA Reverse Transcription Kit. RT-qPCR for target genes (*WTAP*, *METTL3* and *METTL14*) in each fraction were carried out using PowerUp SYBR Master Mix. The target gene mRNA in each fraction was normalized to *GAPDH* in this fraction; the abundance in the heavy polysomes was calculated by summing up fractions #18-25 and compared between UHC and FXS cortical neurons.

### Bulk RNA sequencing and analysis

Total RNAs were extracted from six-week-old cortical neurons using TRIzol reagent followed by mRNAs isolation with the Dynabeads mRNA Purification Kit. Sequencing libraries were prepared using NEBNext Ultra RNA Library Prep kit (New England Biolabs) for Illumina following manufacturer’s protocol. Briefly, mRNAs were fragmented and reverse-transcribed to synthesize double-stranded cDNAs. After end repair and 3’ adenylation, Illumina-compatible adapters for multiplexing were ligated to the ends of the double-stranded cDNA fragments. The ligation products were then amplified and purified to generate sequencing libraries. Sequencing was performed on an Illumina NovaSeq 6000 platform with 150 bp paired end reads at Novogene Bioinformatics Technology Co., Ltd (Beijing, China).

For RNA-seq analysis, similar to the MeRIP-seq workflow, sequence reads obtained from RNA-seq were initially processed and aligned to the human genome (GRCh38) employing STAR (v2.7.9) with default settings. Gene counts were obtained using FeatureCounts (v2.0.3). Differential expression analysis was performed using the R package DESeq2, with a p value cutoff of 0.05 and a fold change threshold of log2 (fold change) > ±1. The differentially expressed genes (DEGs) between FXS and UHC cortical neurons were presented in Table S3. GO enrichment analysis was performed using the clusterProfiler R package (v4.11.0) for functional annotation. CSV files of the DEGs are available in Supplementary table 3. The DESeq2-normalized counts for *METTL3*, *METTL14*, *WTAP*, and *FMR1* transcripts were obtained using variance stabilizing transformation (VST) function in the DESeq2 package.

Statistical comparisons of gene expression were performed following DESeq2 pipeline. Additionally, published MeRIP-seq data for two *Fmr1*-KO and two WT mice (GSE107434) were downloaded from the NCBI Gene Expression Omnibus (GEO) database (https://www.ncbi.nlm.nih.gov/geo/query/acc.cgi?acc=GSE107434) and re-analyzed using the MeRIP-seq input sequencing data, following the aforementioned RNA-seq analysis pipeline for comparing the expression levels of *Mettl3*, *Mettl14*, *Wtap.* The “cor” function in R was used to calculate the Pearson correlation coefficients (PCCs) between DMGs and DEGs.

### RNA stability assay

For six-week-old cortical neurons, treated or untreated with DMSO or STM2457 for 48 hours, gene transcription was inhibited by adding 5 μg/ml of actinomycin D (Cayman Chemical). Cells were harvested at 0-, 2-, and 6-hours post-actinomycin D treatment, washed with PBS, and lysed in TRIzol. Reverse transcription was performed using the High-Capacity cDNA Reverse Transcription Kit. RT-qPCR for *LRRC4B* was then conducted using PowerUp SYBR Master Mix on a QuantStudio system. *LRRC4B* mRNA levels were normalized to *GAPDH*. All primers used are listed in Table S1.

### Data collection and statistics

All experiments were performed in triplicate or more, with data collected from parallel cultures. Quantitative data are presented as mean ± S.E.M. Statistical analyses were conducted using RStudio (v3.4.3) and GraphPad Prism 6. When two independent experimental groups were analyzed, a two-tailed Student t-test was performed. When more than two independent experimental groups were analyzed, one-way ANOVA with Tukey post hoc test was performed. For evoked APs frequency, two-way ANOVA with Tukey post hoc test was performed. Other statistical methods included Kolmogorov-Smirnov test and Pearson’s correlation analysis. A significant threshold of p < 0.05 was applied.

## Data and code availability

The m^6^A-seq data has also been submitted to GEO with accession number GSE278698. The RNA-seq data generated in this study has been deposited in the GEO database under accession number GSE278697. Additional data, including, m^6^A peak calling and annotation, FMRP targets, SAFRI ASD genes, synaptic genes, overlap analyses, and DEGs, are provided in the Supplementary Materials (Tables S2-S4). All other relevant data is available upon request from the corresponding author. Commands used to run software such as STAR, exomePeak, Homer, and Guitar are included in the respective software manuals, which are publicly accessible online. R scripts used for bioinformatics analyses are available upon request.

