## Supplemental Figures for "An iPSC model of fragile X syndrome reflects clinical phenotypes and reveals m^6^A- mediated epi-transcriptomic dysregulation underlying synaptic dysfunction"

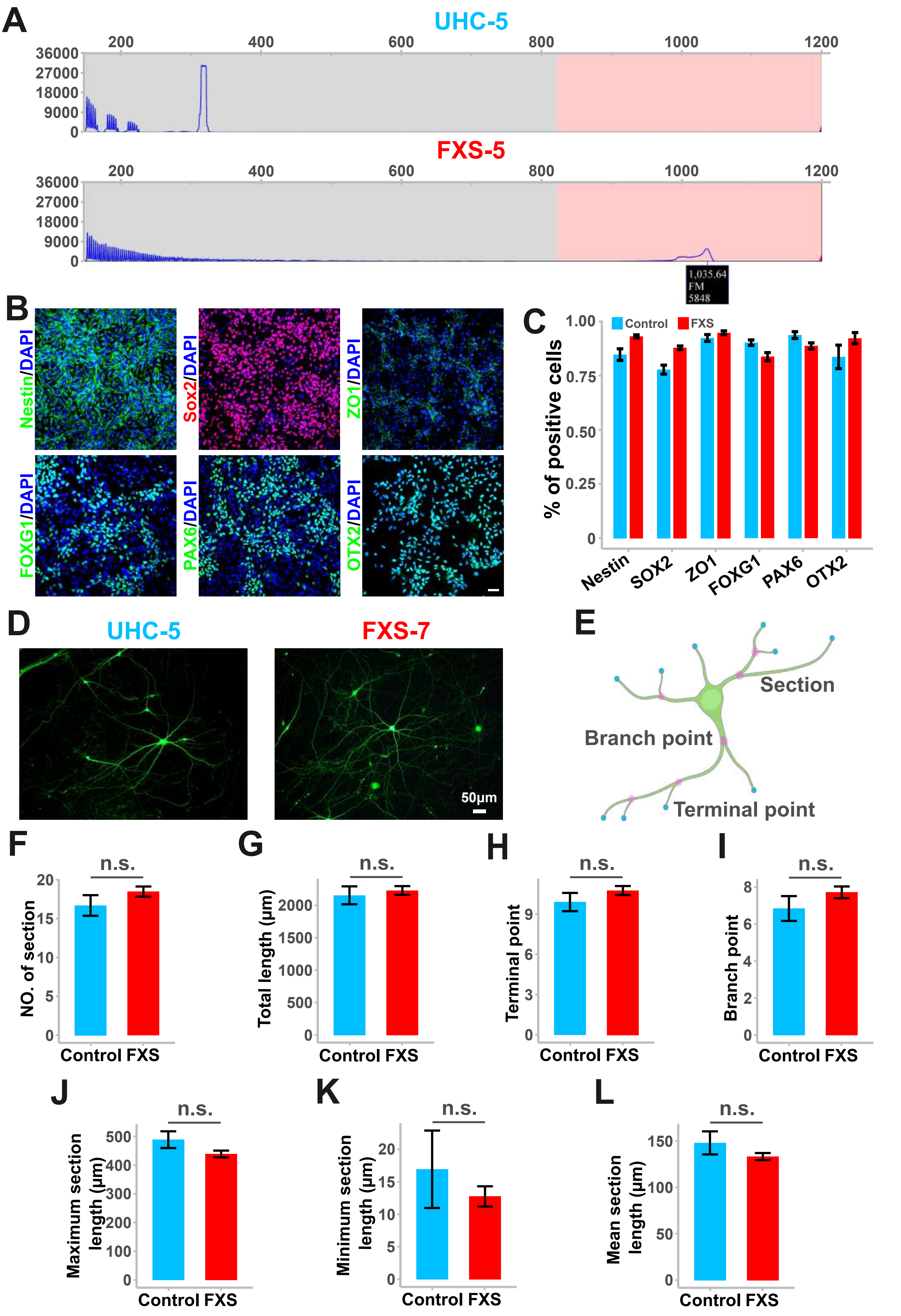


Suppl. Fig 1. (Lu, et al.)

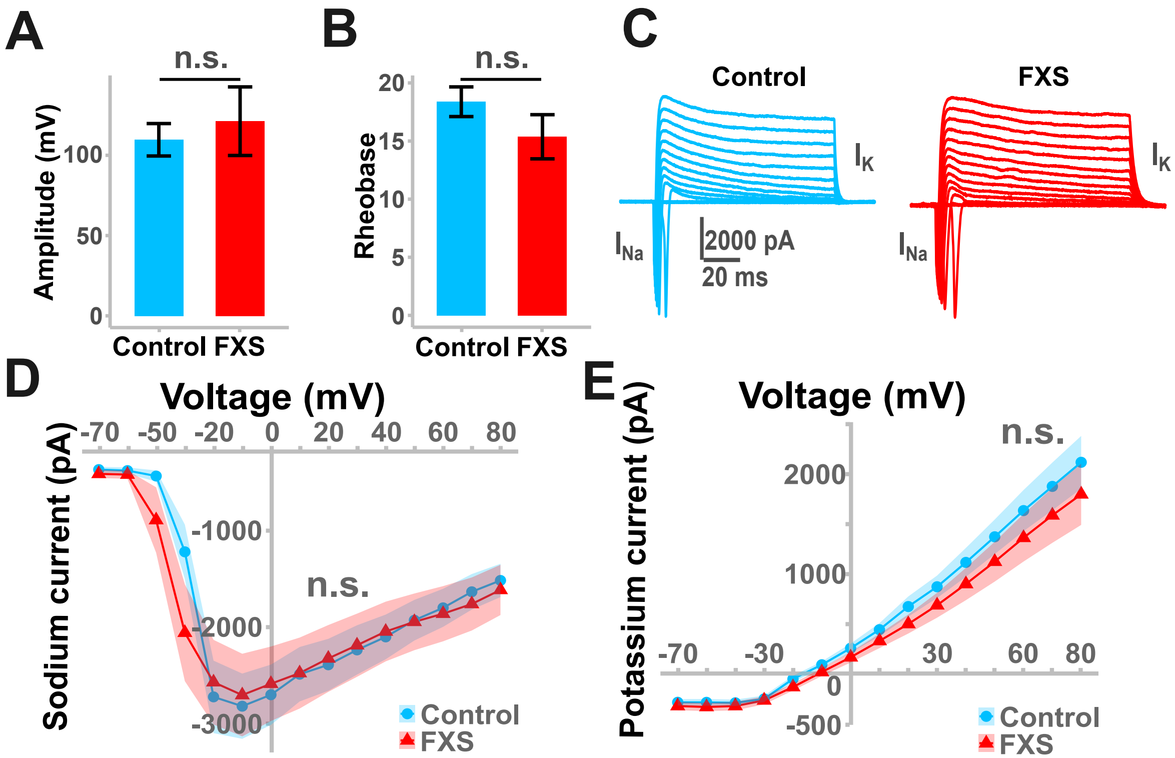


Suppl. Fig 2. (Lu, et al.)

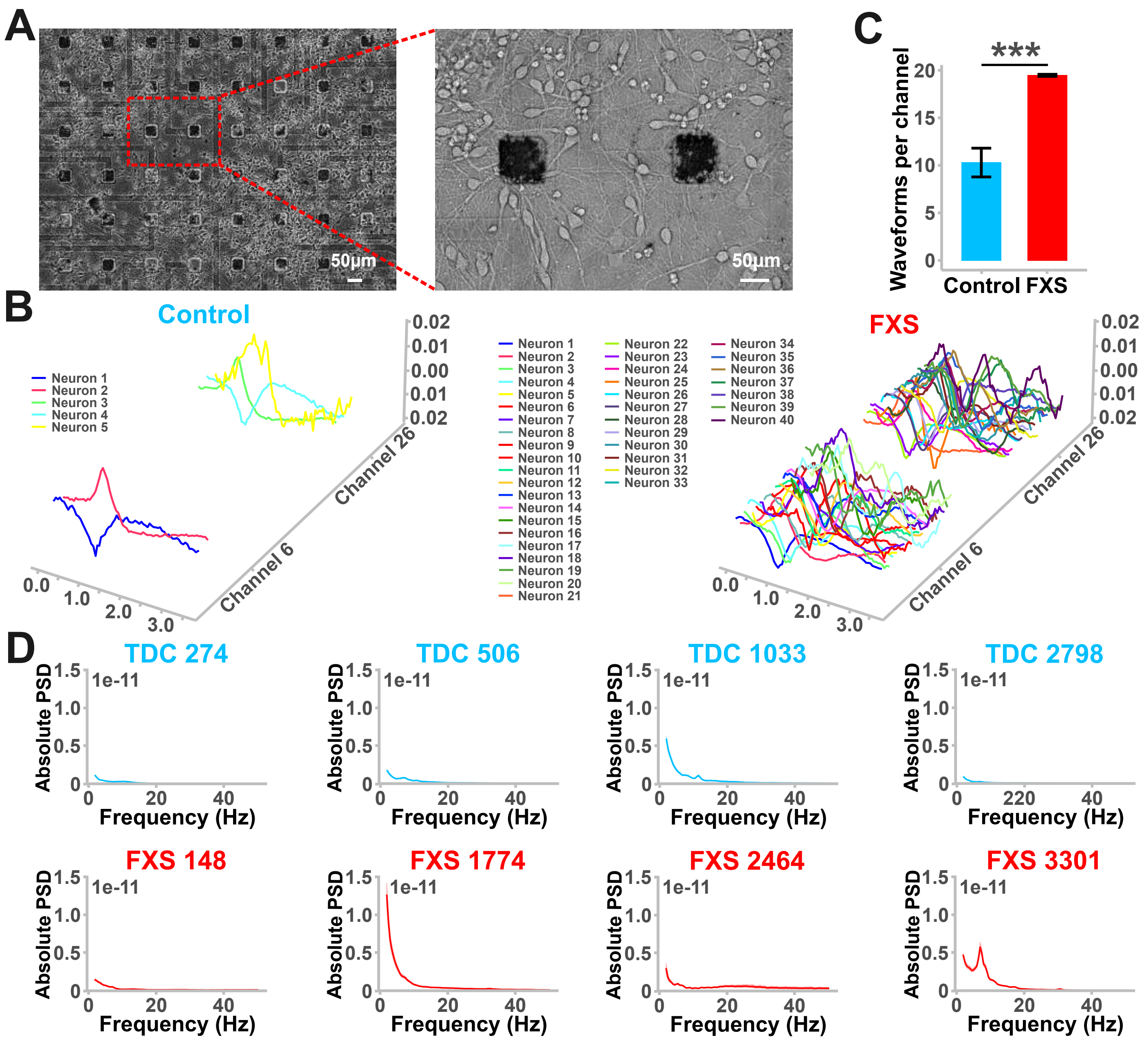


Suppl. Fig 3. (Lu, et al.)

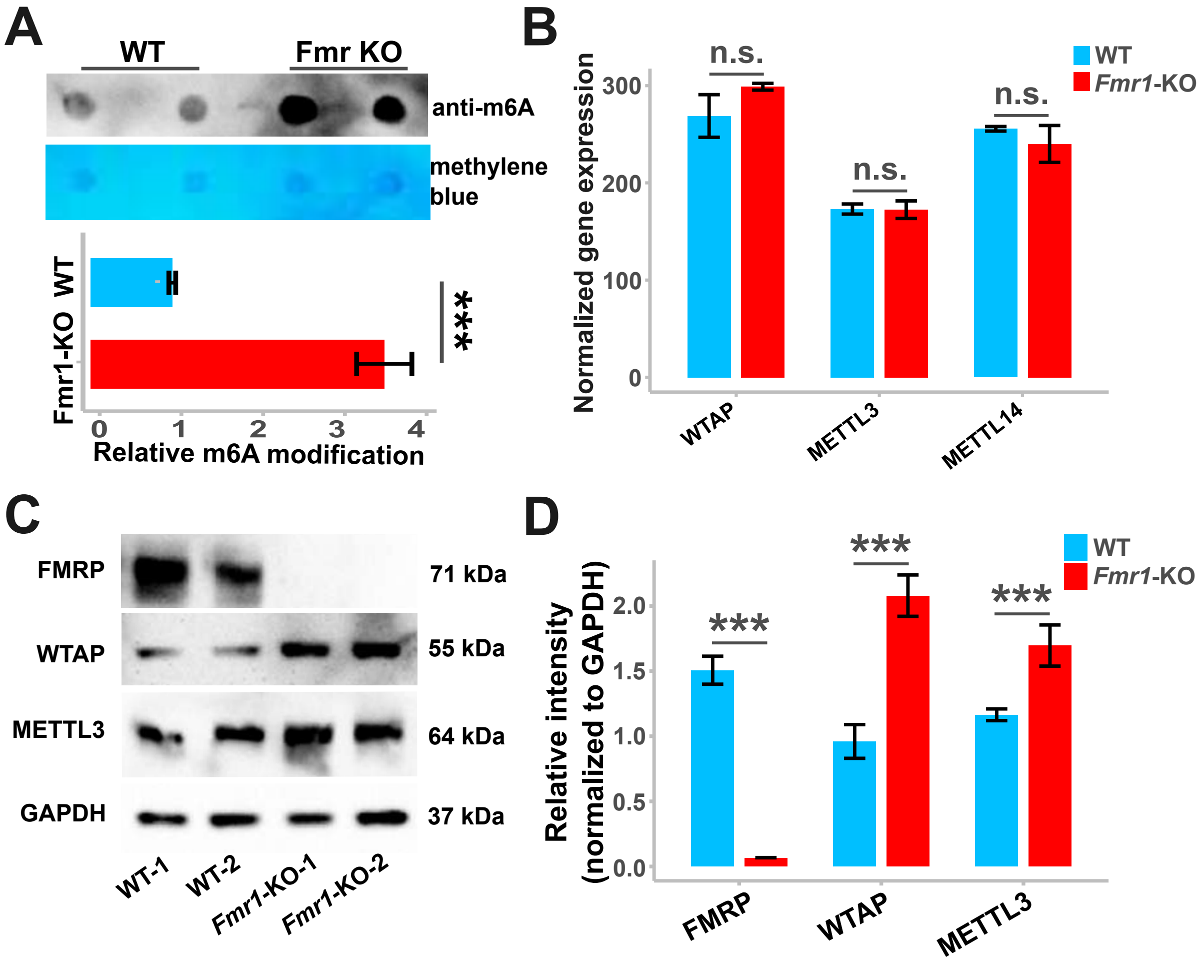


Suppl. Fig 4. (Lu, et al.)

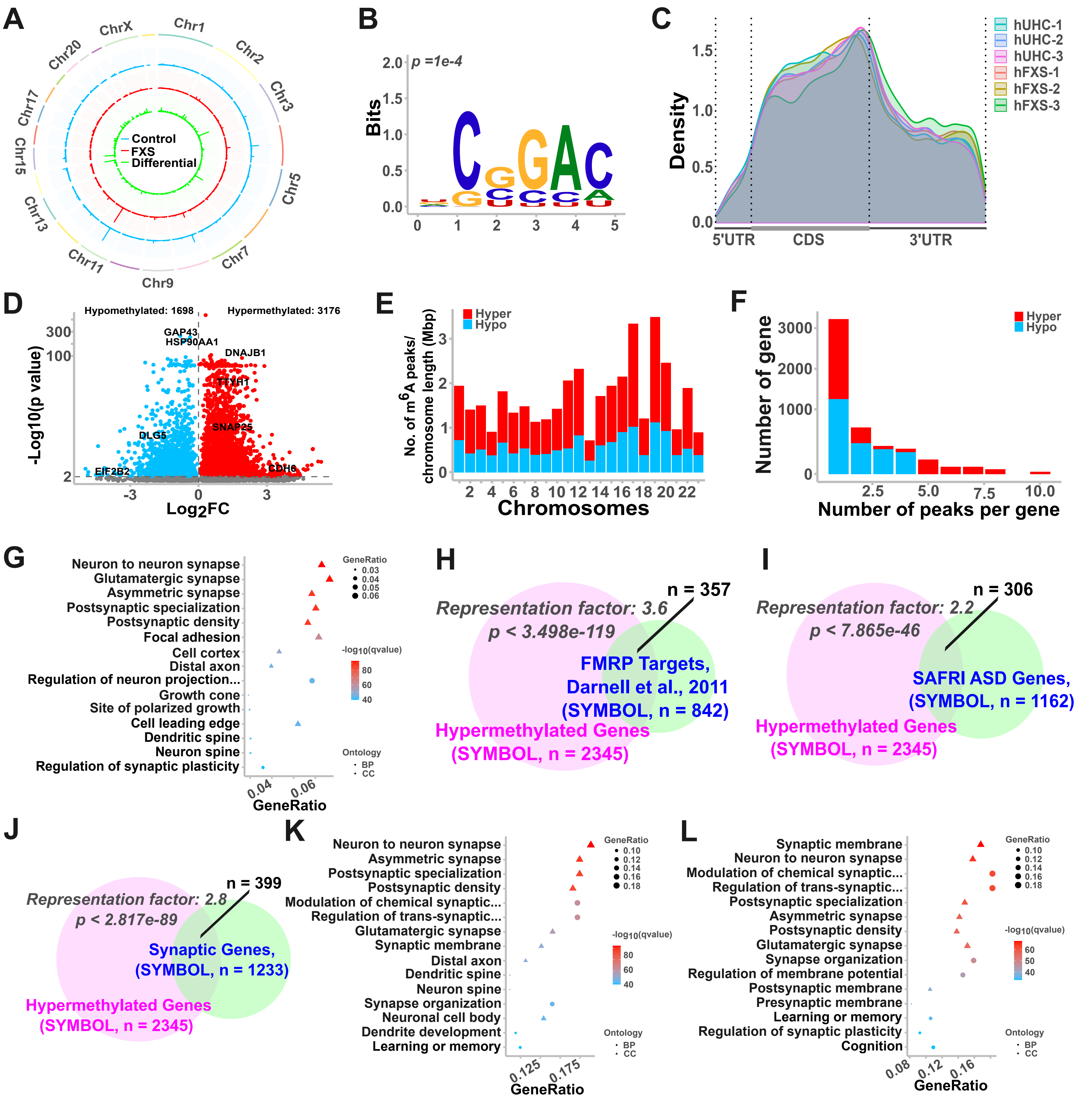


Suppl. Fig 5. (Lu, et al.)

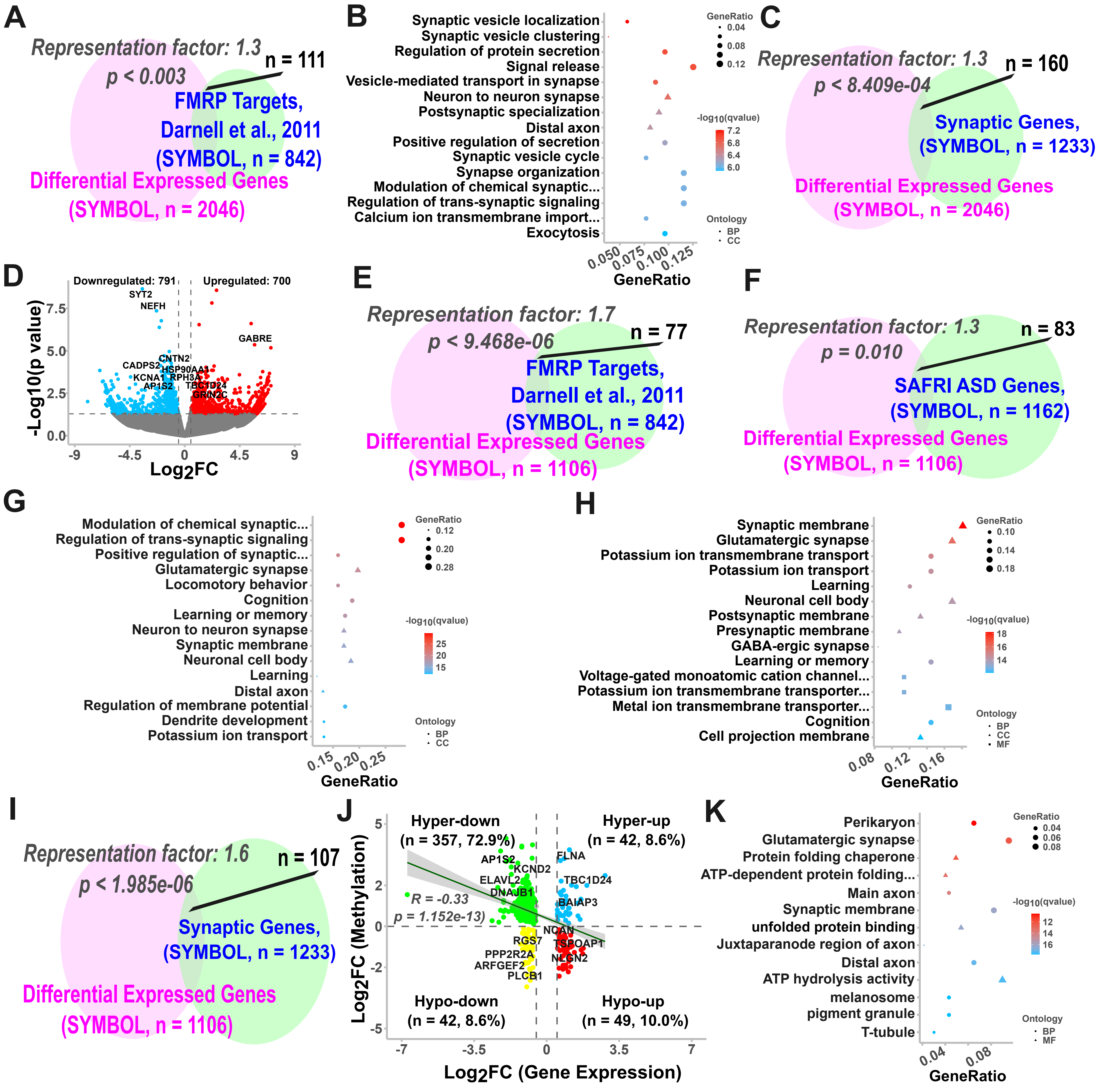


Suppl. Fig 6. (Lu, et al.)
